# The robust, high-throughput, and temporally regulated roxCre and loxCre reporting systems for genetic modifications *in vivo*

**DOI:** 10.1101/2024.04.23.590680

**Authors:** Mengyang Shi, Jie Li, Xiuxiu Liu, Kuo Liu, Lingjuan He, Wenjuan Pu, Wendong Weng, Shaohua Zhang, Huan Zhao, Kathy O. Lui, Bin Zhou

**Affiliations:** CAS CEMCS-CUHK Joint Laboratories, New Cornerstone Science Laboratory, State Key Laboratory of Cell Biology, CAS Center for Excellence in Molecular Cell Science, Shanghai Institute of Biochemistry and Cell Biology, Chinese Academy of Sciences, University of Chinese Academy of Sciences, Shanghai, China; Key Laboratory of Systems Health Science of Zhejiang Province, School of Life Science, Hangzhou Institute for Advanced Study, University of Chinese Academy of Sciences, Hangzhou, China; School of Life Sciences, Westlake University, Hangzhou, China; Department of Chemical Pathology, Li Ka Shing Institute of Health Sciences, The Chinese University of Hong Kong, Prince of Wales Hospital, Hong Kong, China; School of Life Science and Technology, ShanghaiTech University, Shanghai, China

## Abstract

Cre-loxP technology, a cornerstone in fate mapping and *in vivo* gene function studies, faces challenges in achieving precise and efficient conditional mutagenesis through inducible systems. This study introduces two innovative genetic tools designed to overcome these limitations. The first, roxCre, enables DreER-mediated Cre release, paving the way for intersectional genetic manipulation that permits increased precision and efficiency. The second, loxCre, facilitates conditional gene targeting by allowing CreER lines to induce Cre expression with significantly enhanced efficiency. These tools incorporate a fluorescent reporter for genetic lineage tracing, simultaneously revealing efficient gene knockout in cells marked by the reporter. These strategies hold great potential for precise and efficient exploration of lineage-specific gene functions, marking a significant advancement in genetic research methodologies.

## Introduction

Genetic modification, such as gene knockout in a precise spatiotemporal manner forms the basis for understanding cellular and molecular mechanisms in multiple biological processes^1, 2^. The Cre-loxP recombination system is widely employed to knock out or over-express specific genes *in vivo*^3^. Cre recombinase, derived from the bacteriophage P1 gene, targets short palindrome DNA sequence loxP (34bp) and recombines codirectional-loxP-flanked sequences for deletion^4, 5^. To date, most of the mouse genes have been successfully targeted by two loxP sites (floxed allele), which could be recombined by cell-type specific Cre lines. Application of the Cre-loxP system in mouse genetics revolutionizes gene functional analysis and significantly advances our exploration of many developmental and pathophysiological processes at the molecular level^6, 7^.

While constitutively active Cre is robust and efficient in recombination for gene deletion, Cre lacks temporal control as the promoter driving Cre is active from the embryonic to the adult stage. Therefore, a temporally active Cre is needed to avoid early mortality and realize the ablation of an indispensable gene at a key developmental stage. To overcome the temporal limitation of Cre, CreER, the fusion of Cre with a mutated estrogen receptor, is generated to enable its activation only after tamoxifen (Tam) treatment, and the interaction of ER with Tam induces nuclear translocation of CreER from the cytoplasm^8^. As a result, Tam treatment restricts CreER activity at a specific time window and permits gene functional analysis at a higher spatiotemporal resolution than Cre^9^. As the loxP-flanked sequences and their locations in genome or chromatin influence its accessibility for recombination, the recombination efficiency of CreER varies significantly among multiple floxed alleles, ranging from easy- to inert-to-recombine alleles^10^. For example, CreER has high efficiency in targeting the easy-to-recombine alleles, such as generic reporter *R26-tdT*^11^, but exhibits low efficiency on some other reporters such as *R26-Confetti*^12^, or may not excise some floxed gene alleles that are inert or resistant for targeting, leading to failure of gene deletion in some tdT^+^ cells and causing false-positive tracing fate (Figure 1A). In other words, tdT expression may not always necessarily denote gene deletion in the labeled cells. The notion that the pattern of Cre-expression based on a reporter (e.g., *R26-tdT*) represents the pattern of Cre-mediated recombination is less rigorous^13^. Therefore, it is not precise to assume the successful deletion of codirectional-loxP-flanked sequences in the gene of interest based on the deletion in another gene (or reporter) by the same CreER mouse line.

**Figure 1.**
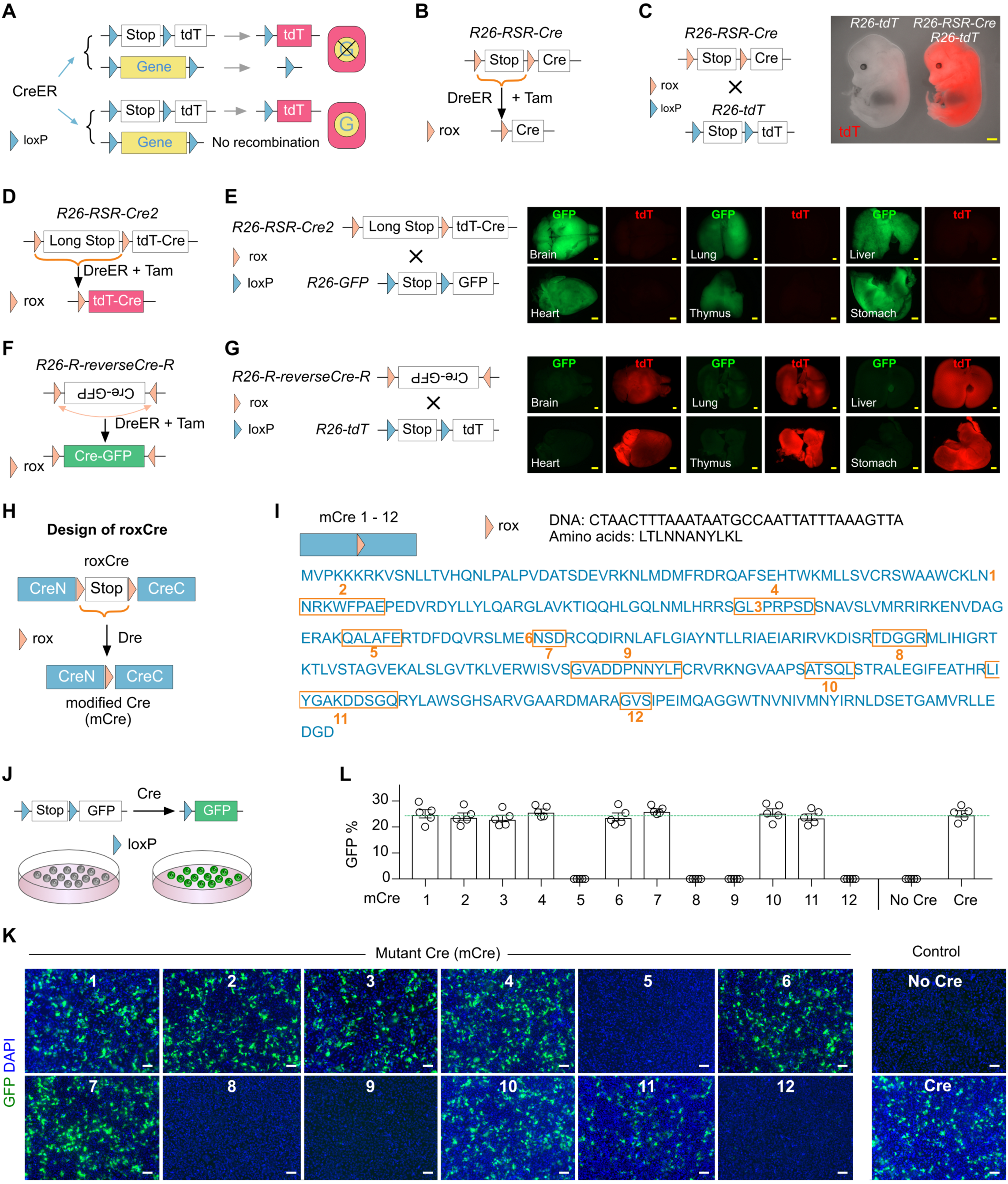
Design of roxCre for DreER-induced mCre expression. (A) A schematic showing genetic labeling and/or gene knockout in CreER-expressing cells. (B, D, F) Strategies for DreER-induced Cre expression. (C) Crossing with *R26-tdT* mice, *R26-RSR-Cre* exhibits leakiness in E13.5 embryos. Scale bars, yellow 1mm. Each image is representative of 5 individual biological samples. (E) Examination of Cre leakiness by whole-mount fluorescence images of organs collected from *R26-RSR-Cre2;R26-GFP* adult mice. Scale bars, yellow 1mm. Each image is representative of 5 individual biological samples. (G) Examination of Cre leakiness by whole-mount fluorescence images of organs collected from *R26-R-reverseCre-R;R26-tdT* adult mice. Scale bars, yellow 1mm. Each image is representative of 5 individual biological samples. (H) A schematic showing the design for roxCre and mCre. (I) A schematic showing mCre1 to mCre12 by insertion of rox into *Cre* cDNA at 12 different loci. (J) Examination of mCre for recombination efficiency by the *loxP-Stop-loxP-GFP* reporter. (K) Immunofluorescence images of cells stained with GFP on cells. Scale bars, white, 100 µm. Each image is representative of 5 individual biological samples. (L) Quantification of the percentage of cells expressing GFP in each group. Data are the means ± SEM; n = 5.

One way to increase the recombination efficiency of CreER is to treat mice with many times of Tam, as the high dosage of Tam in theory could increase the chances of CreER-mediated recombination. However, the increased dosage of Tam treatment is toxic and could lead to many side effects on the phenotype, creating confounding effects on the study^14, 15^. To increase the inducible recombination efficiency of floxed alleles, Lao et al. reported a mosaic mutant analysis with spatial and temporal control of recombination (MASTR), in which initial FlpoER-mediated recombination induces GFPCre expression, that subsequently recombines floxed alleles effectively in GFP^+^ cells^16^. Similarly, a Flp-induced mosaic analysis system with Cre or Tomato (MASCOT) has been reported to express Cre and reporter simultaneously in a cell^17^. While MASTR and MASCOT enable spatiotemporal labeling of mosaic mutant cells utilizing the current resources of floxed mouse lines, it may not be suitable for effective gene knockout at a tissue/population level due to low recombination efficiency initiated by Flp recombinase. In order to improve the inducible recombination efficiency, Tian et al. have also reported a self-cleaved inducible CreER (sCreER) that switches inducible CreER into a constitutively active Cre, effectively recombining floxed allele for gene manipulation^10^. However, self-cleaved inducible Cre mice have to be newly generated each time for targeting different promoters, which is time-consuming and the system is not compatible with current resources largely based on CreER mouse lines. A recent study has reported an iSuRe-Cre strategy to induce and report Cre-dependent genetic modifications utilizing conventional CreER tools^18^. The reporter expression reflects gene knockout in cells by iSuRe-Cre. However, two limitations hinder its widespread applications. First, without any induction, the iSuRe-Cre transgene is leaky in some tissues, such as heart and skeletal muscle. Second, the initial recombination for releasing the iSuRe-Cre allele induced by CreER is far from efficient in some tissues compared with easy-to-recombine alleles such as *R26-tdT*^18^, limiting its usage for efficient gene knockout at a tissue level. Thus, a new method is needed to ensure gene knockout in cells as effectively and efficiently as recombination on easy-to-recombine alleles, thus allowing efficient gene deletion at a population level.

Conventional Cre-loxP system uses tissue-specific promoter to drive Cre, thus the precision of genetic targeting solely depends on promoter activity. Now it is known that many promoters are not as specific as previously reported^19–21^. The ectopic or unwanted Cre expression in other cell types may result in unspecific cell labeling or gene deletion, leading to many confounding issues or controversies in multiple fields of research^22–24^. In addition, some cell types cannot be clearly distinguished by one marker from another, and thus could not be specifically and genetically targeted by only one gene promoter-driven recombinase^25^. To circumvent this limit, dual recombinases-mediated genetic targeting utilizes two promoters to independently drive Cre and Dre^26^. Dre is another bacteriophage recombinase that targets rox sites and is orthogonal to the Cre-loxP system^27^. While cells of interest could be labeled by specific reporters that are responsive to dual recombinases, gene deletion requires a final readout on singular Cre recombinase for targeting floxed gene alleles.

In this study, we developed two genetic strategies: roxCre and loxCre, in which the *rox-Stop-rox* (RSR) and *loxP-Stop-loxP* cassettes are respectively inserted into the *Cre* coding region, such that Cre transcription and translation are not properly carried out before the removal of the *Stop* cassette. There is no obviously spontaneous leakiness by roxCre or loxCre. We found that CreER-induced loxCre effectively deletes genes in reporter-labeled cells, ensuring efficient gene knockout for the evaluation of lineage-specific gene function. In addition, DreER-induced roxCre efficiently deletes genes in cells for both specific and efficient intersectional genetic studies. We expect that these tools would overcome the issues of inefficient and nonspecific genetic modifications, enhancing the ability to more precisely manipulate genes in a particular cell lineage for a better understanding of gene functions in multiple biomedical fields.

## Results

### Design of roxCre for DreER-induced mCre expression

A sequential genetic approach has been recently reported to delete genes^28^, which used Dre-induced CreER expression by removing the *rox-Stop-rox* sequence ahead. As aforementioned, the recombination carried out by the released CreER for excising floxed gene alleles may not be as efficient as reporters (e.g., *R26-tdT*). A possible strategy to control Cre activity is to engineer a rox-flanked *Stop* sequence ahead of its DNA (*R26-RSR-Cre*) that will be removed after Dre-rox recombination (Figure 1B). However, such a strategy is not ideal, as the upstream transcriptional *Stop* sequence might not completely prevent *Cre* transcription, leading to leakiness (Figure 1C). We then inserted a longer *Stop* sequence or reversed the direction of the *Cre* DNA by generating *R26-RSR-Cre2* and *R26-R-reverseCre-R* lines, respectively. The whole-mount fluorescence imaging results displayed heavy leakiness, which demonstrates that Cre was independently activated for recombination without being crossed with a Dre mouse line (Figure 1D-G). To enable efficient dual recombinases-mediated gene knockout, we need to design a strategy for DreER-induced expression of a constitutively active Cre, and the system should not exhibit leakiness.

We first generated roxCre, in which an RSR cassette was inserted into the coding region of *Cre*, splitting *Cre* into N- and C-terminal segments (Figure 1H). After removal of RSR by Dre-rox recombination, Cre is recombined containing one remaining rox site within its coding sequence, hereafter termed modified Cre (mCre, Figure 1H). To ensure that mCre has the same recombination efficiency as the conventional Cre, we inserted a rox sequence at 12 different sites of *Cre* individually, most of which were located between helix domains (Figure 1I). As insertions of additional amino acids of rox would change the original sequence of Cre, potentially leading to reduced activity, we screened for their recombination efficiency by transfecting cells with 12 versions of mCre (mCre1 to mCre12) and the responding GFP reporter (Figure 1J). We found that mCre1, 2, 3, 4, 6, 7, 10, and 11 displayed a comparable efficiency as the conventional Cre; whereas mCre5, 8, 9, and 12 exhibited impaired recombination efficiency (Figure 1K, L). These data demonstrate that some of the mCre generated in our study are as efficient as the conventional Cre *in vitro*.

Next, we sought to screen for the most robust version of mCre *in vivo.* We generated four roxCre knock-in mice based on mCre1, 4, 7, and 10 as determined from the above *in vitro* screening (Figure S1-S2). In these roxCre lines, RSR inserted into *Cre* was removed by Dre-rox recombination resulting in the generation of mCre (Figure 1J). We then used hepatocyte- and endothelial cell-specific promoters albumin (Alb) and VE-cadherin (Cdh5) to drive roxCre, and generate *Alb-roxCre1-tdT*, *Cdh5-roxCre4-tdT*, *Alb-roxCre7-GFP*, and *Cdh5-roxCre10-GFP* knock-in mice, respectively (Figure S1-S2). No spontaneous fluorescence signal was detected in *Alb-roxCre1-tdT*, *Cdh5-roxCre4-tdT*, *Alb-roxCre7-GFP*, and *Cdh5-roxCre10-GFP* knock-in mice (Figure S1-S2), demonstrating no reporter leakiness without Dre-rox recombination. Nor did we detect leakiness as a result of any Cre activity when they were crossed with *R26-GFP*^29^ or *R26-tdT*^11^ reporter (Figure S1-S2). These data demonstrate that roxCre mouse lines are functionally efficient yet non-leaky.

### DreER-induced mCre robustly recombines inert alleles

To more rigorously evaluate the recombination effectiveness among different versions of mCre, we used an inducible DreER to temporally and specifically release mCre in hepatocytes or endothelial cells. Generic reporter mouse lines *R26-GFP* and *R26-tdT* were used to indicate the difference in recombination efficiency between roxCre1 and roxCre7 first. The fluorescence-activated cell sorting (FACS) and immunostaining results showed no leakiness in *R26-DreER;Alb-roxCre1-tdT;R26-GFP* and *R26-DreER;Alb-roxCre7-GFP;R26-tdT* (Figure S3C-F). Nevertheless, both two versions of mCre efficiently labeled all targeted cells (Figure S3C-E, G) due to the easy-to-recombine characteristic of *R26-GFP* and *R26-tdT*.

To evaluate a strong mCre, we attempted to target one of the most inert alleles for recombination, *R26-Confetti*^12^, which is often used as reporters for clonal analysis due to its sparse labeling with rare recombination events^10, 12^. According to the Cre-loxP recombination principle, Cre can remove the sequence between two loxP sites that have the same transcription direction and turn over the sequence in the middle of two loxP sites that have the opposite transcription direction. As the recombination efficiency of CreER is limited, *R26-Confetti* can only be recombined into YFP, or nGFP (nuclear GFP), or mCFP (membrane CFP), or RFP (Figure S4A).

We crossed *R26-DreER;R26-Confetti* mice with *Alb-roxCre1-tdT* (group 1) and *Alb-roxCre7-GFP* (group 2) mice, and Tam-induced Dre-rox recombination yielded *Alb-mCre1-tdT* and *Alb-mCre7-GFP* mice, respectively (Figure 2A). In this system, the constitutively active mCre would completely remove the sequence between two loxP sites that have the same transcription direction and recombine *R26-Confetti* into two sets of reporter pairs: YFP-nGFP pair and mCFP-RFP pair, which could be detected in a single hepatocyte due to the poly-nuclear or polyploidy feature of hepatocytes (Figure 2A). In each pair, reporters can equally express, as mCre is strong enough to turn over the sequence in the middle of two loxP sites that have the opposite transcription direction (Figure S4B). Therefore, we used YFP/mCFP to detect mCre recombination efficiency and tdT/RFP or GFP to trace its endogenous reporter activity that was driven by Alb promoter in *Alb-mCre1-tdT* and *Alb-mCre7-GFP* mice, respectively (Figure 2A). Besides, *Alb-CreER*^26^*;R26-Confetti* was used as a control under the same Tam treatment (group 3, Figure 2A).

**Figure 2.**
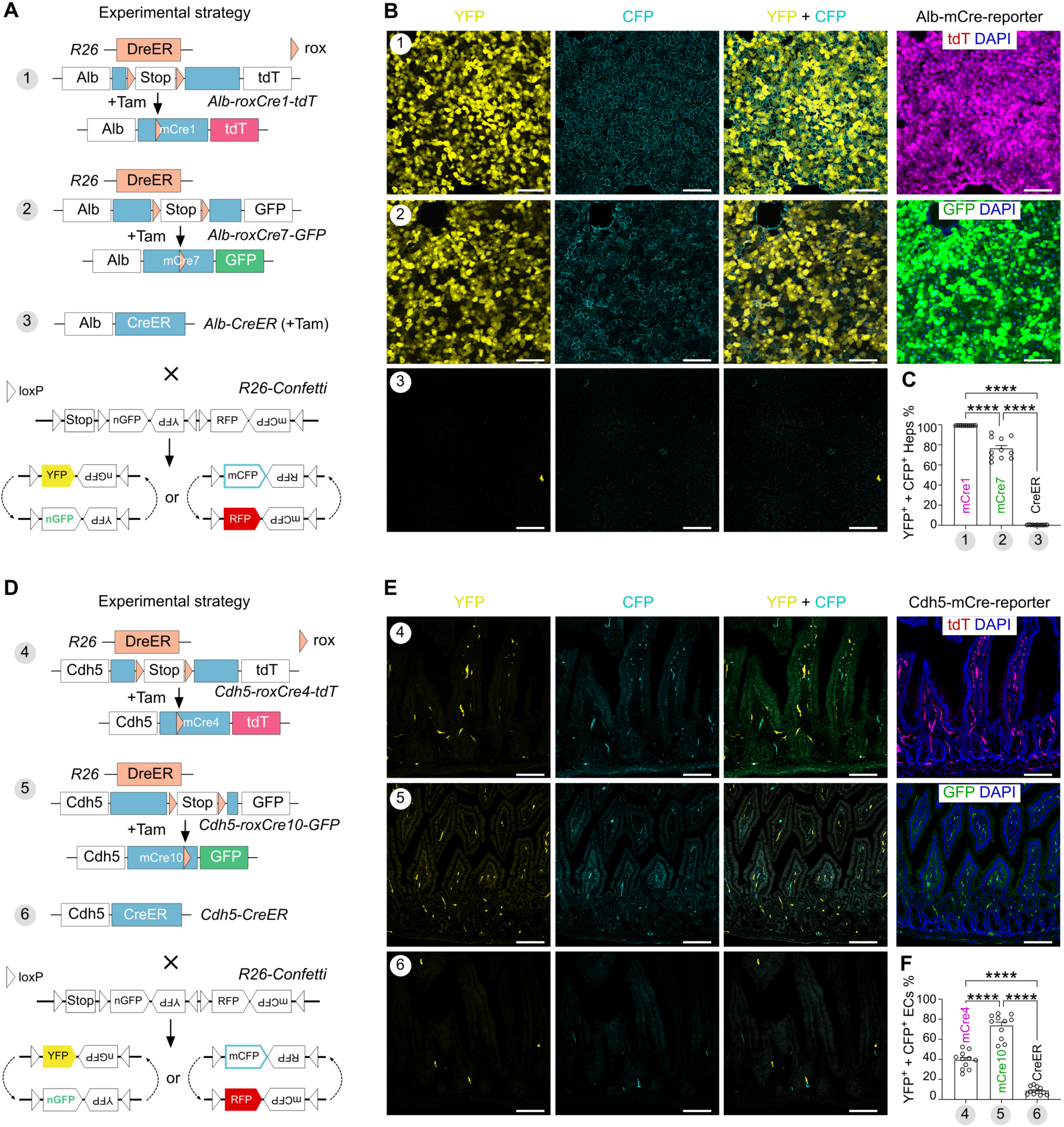
DreER-induced mCre robustly recombines inert alleles. (A) A schematic showing the experimental design to test the recombination efficiency of mCre1 and mCre7 on the *R26-Confetti* allele. Tam-induced DreER-rox recombination leads to mCre1/tdT or mCre7/GFP expression in hepatocytes in strategy 1 or 2, respectively. Strategy 3 uses conventional *Alb-CreER* as control. YFP and mCFP signals are used for the examination of recombination on the *R26-Confetti* allele. (B) Fluorescence images of YFP and mCFP on liver sections collected from mice in strategy 1-3. Strategy 1 and 3 exhibit tdT or GFP in hepatocytes after DreER-rox recombination, respectively. Scale bars, 100 µm. Each image is representative of 11 individual biological samples. (C) Quantification of the percentage of hepatocytes (Heps) expressing either YFP and/or mCFP in strategy 4-6. Data are the means ± SEM; n = 11 mice for each strategy; \*\*\*\**P* < 0.0001 by one-way ANOVA. (D) A schematic showing the experimental strategy to test the recombination efficiency of mCre4 and mCre10 on *R26-Confetti* allele. Tam-induced recombination leads to mCre4/tdT or mCre10/GFP expression in endothelial cells (ECs) in strategy 4 or 5, respectively. Strategy 6 uses conventional *Cdh5-CreER* as control. YFP and mCFP signals are used for the examination of recombination on *R26-Confetti* allele. (E) Fluorescence images of YFP and mCFP on intestinal sections collected from mice in strategy 4-6. Strategy 4 and 5 exhibit tdT or GFP in ECs after DreER-rox recombination, respectively. Scale bars, 100 µm. Each image is representative of 11 individual biological samples. (F) Quantification of the percentage of ECs expressing either YFP and/or mCFP in strategy 4-6. Data are the means ± SEM; n = 11 mice for each strategy; \*\*\*\**P* < 0.0001 by one-way ANOVA.

In fact, DreER-rox recombination in hepatocytes is not 100%. Therefore, we quantified YFP and mCFP expression in hepatocytes with DreER-rox recombination to evaluate the strength of mCre activity. We found sparse YFP^+^ or mCFP^+^ hepatocytes (0.39 ± 0.04%) in *Alb-CreER;R26-Confetti* mice. However, 99.94 ± 0.02% of tdT^+^ hepatocytes (DreER recombined) were YFP^+^ or mCFP^+^ in *R26-DreER;Alb-roxCre1-tdT;R26-Confetti* mice; and 76.83 ± 3.20% of GFP^+^ hepatocytes (DreER recombined) were YFP^+^ or mCFP^+^ in *R26-DreER;Alb-roxCre7-GFP;R26-Confetti* mice (Figure 2B, C). Similarly, we found that 39.81 ± 2.61% of tdT^+^ endothelial cells and 74.15 ± 3.64% of GFP^+^ endothelial cells in small intestine were YFP^+^ or mCFP^+^ in *R26-DreER;Cdh5-roxCre4-tdT;R26-Confetti* and *R26-DreER;Cdh5-roxCre10-GFP;R26-Confetti* mice, respectively, compared with sparse labeling in *Cdh5-CreER;R26-Confetti* mice (Figure 2D-F). Similar results were also observed in other tissues and organs (Figure S5). Furthermore, *R26-DreER;Alb-roxCre1-tdT;R26-Confetti*, *R26-DreER;Alb-roxCre7-GFP;R26-Confetti*, *R26-DreER;Cdh5-roxCre4-tdT;R26-Confetti*, *and R26-DreER;Cdh5-roxCre10-GFP;R26-Confetti* were not leaky (Figure S4C). Together, these data demonstrate that inducible DreER controls roxCre activation, and mCre1 is the most efficient recombinase assisting the recombination of inert alleles *in vivo*. Hereafter, roxCre1 will be referred to as roxCre for brevity.

### Dre-induced mCre realizing cell subpopulation-specific gene manipulation

To substantiate the precision and specificity of roxCre targeting cells, we employed intersectional genetic targeting using dual recombinases. We generated the *Cyp2e1-DreER* mouse line which labeled peri-central hepatocytes and also renal epithelial cells (Figure S6). Crossing *Cyp2e1-DreER* with hepatocyte-specific *Alb-roxCre-tdT* mice achieved genetic targeting of peri-central hepatocytes exclusively, circumventing issues associated with ectopic targeting of renal cells by *Cyp2e1-DreER* line (Figure S6) or unwanted targeting of peri-portal hepatocytes by *Alb-CreER* line^26^. Wnt-β-catenin plays an important role in liver development, zonation, homeostasis, and diseases^30^. Therefore, we generated the *Alb-roxCre-tdT;Ctnnb1^flox/flox^* mouse line to examine if Dre-induced mCre could efficiently delete *Ctnnb1* gene that encodes β-catenin (*Ctnnb1^flox^*)^31^.

We generated *Cyp2e1-DreER;Alb-roxCre-tdT;Ctnnb1^flox/flox^*triple knock-in mice (mutant), in which Tam-induced DreER-rox recombination yielded mCre that subsequently targeted floxed *Ctnnb1* allele (Figure 3A). We treated mutant mice and their littermate control *Cyp2e1-DreER;Alb-roxCre-tdT;Ctnnb1^flox/+^* mice with Tam at 8 weeks of age, and sorted tdT^+^ hepatocytes for analysis at 3 days after Tam treatment (Figure 3B). qRT-PCR analysis showed significantly reduced *Ctnnb1* and *Glul* in tdT^+^ hepatocytes of the mutant group, compared to that of the control group (Figure 3C, D). We then collected liver samples at 3 days (D) and 4 weeks (W) post-Tam for further analysis (Figure 3E). Immunostaining for tdT, β-catenin, GS, and E-CAD on liver sections revealed similar levels of β-catenin and GS expression in the 3D mutant liver compared to the control liver (Figure 3F). However, expression of β-catenin and GS reduced in the 4W mutant liver tdT^+^ region compared with the control (Figure 3G), suggesting efficient deletion of *Ctnnb1* that participates in regulating Wnt signaling downstream gene target *Glul*, which encodes GS. Western blotting against β-catenin and qRT-PCR against *Ctnnb1* of sorted tdT^+^ hepatocytes at 4W post-Tam revealed the significant *Ctnnb1* knock-out (Figure 3H-K). Additionally, Wnt downstream target genes including *Glul*, *Axin2*, *Cyp1a2*, *Cyp2e1*, *Oat*, *Tcf7*, *Lect2*, *Tbx3*, *Slc1a2*, and *Rhbg* were significantly reduced in the mutant sorted tdT^+^ hepatocytes than the control group (Figure 3L). Taken together, these data demonstrate that DreER-induced mCre enables efficient intersectional genetic manipulation in the cells of interest.

**Figure 3.**
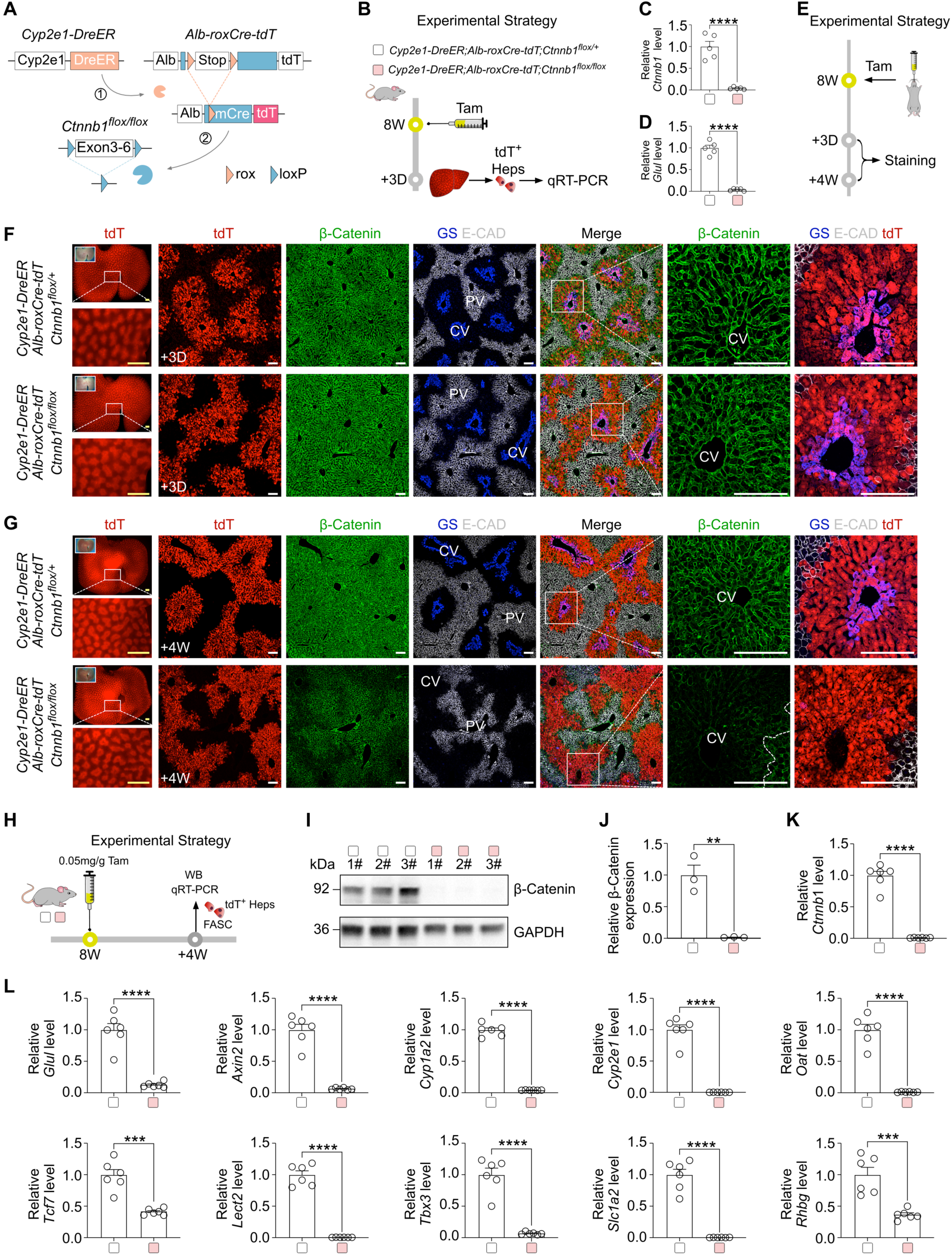
Dre-induced mCre efficiently deletes genes in specific cell subpopulation. (A) A schematic showing the experimental design for Dre-induced mCre expression and the subsequent gene deletion. (B) A schematic showing the experimental strategy. (C and D) qRT-PCR analysis of the relative expression of *Ctnnb1* (C) and *Glul* (D) in the sorted tdT^+^ hepatocytes. Data are the means ± SEM; n = 5; \*\*\*\**P* < 0.0001 by student’s *t* test. (E) A schematic showing the experimental strategy. (F) Immunostaining for tdT, β-Catenin, GS, and E-CAD on liver sections collected on the day 3 post-Tam. PV, portal vein; CV, central vein. Scale bars, yellow 1mm; white 100 µm. Each image is representative of 5 individual biological samples. (G) Immunostaining for tdT, β-Catenin, GS, and E-CAD on liver sections collected at week 4 post-Tam. Scale bars, yellow 1mm; white 100 µm. Each image is representative of 5 individual biological samples. (H) A schematic showing the experimental strategy. (I) Western blotting of β-Catenin and GAPDH in sorted tdT^+^ cells. (J) Quantification of the relative expression of β-Catenin protein. Data are the means ± SEM; n = 3. \*\**P* < 0.01 by student’s *t* test. (K) qRT-PCR analysis of the relative expression of *Ctnnb1* in sorted tdT^+^ cells. Data are the means ± SEM; n = 6. \*\*\*\**P* < 0.0001 by student’s *t* test. (L) qRT-PCR analysis of the relative expression of *Glul*, *Axin2*, *Cyp1a2*, *Cyp2e1*, *Oat*, *Tcf7*, *Lect2*, *Tbx3*, *Slc1a2,* and *Rhbg* in sorted tdT^+^ cells. Data are the means ± SEM; n = 6. \*\*\**P* < 0.001, \*\*\*\**P* < 0.0001 by student’s *t* test.

### *R26-loxCre-tdT* is efficiently recombined by CreER

Having successfully constructed roxCre for Dre-induced mCre expression, we iterated the system to enable mCre induction by CreER, a more widely available tool for biomedical research. We replaced the rox sites of roxCre1 with loxP sites by inserting a *loxP-Stop-loxP* cassette into *Cre* sequence at the same locus as roxCre1(Figure 4A). We then generated the *R26-loxCre-tdT* mouse line, in which mCre and tdT would be expressed simultaneously after *Stop* removal, allowing tdT as a surrogate marker for mCre detection in the same cell (Figure 4C). We then designed two experiments to verify the baseline leakiness of Cre expression (Figure 4B) and to examine if mCre/tdT could be expressed after *Stop* removal (Figure 4C), respectively, by *R26-loxCre-tdT;R26-tdT* (Strategy 1) and by injecting AAV8-Cre virus into *R26-loxCre-tdT* mice (Strategy 2). Immunostaining for tdT on tissue sections revealed no tdT expression in Strategy 1, demonstrating minimal to negligible leakiness at the baseline. The robust tdT expression in the liver in Strategy 2 showed mCre/tdT was allowed to express after *Stop* removal by AAV2/8-hTBG-Cre (Figure 4D). The FACS results confirmed that no leakiness in *R26-loxCre-tdT* (Figure S7A).

**Figure 4.**
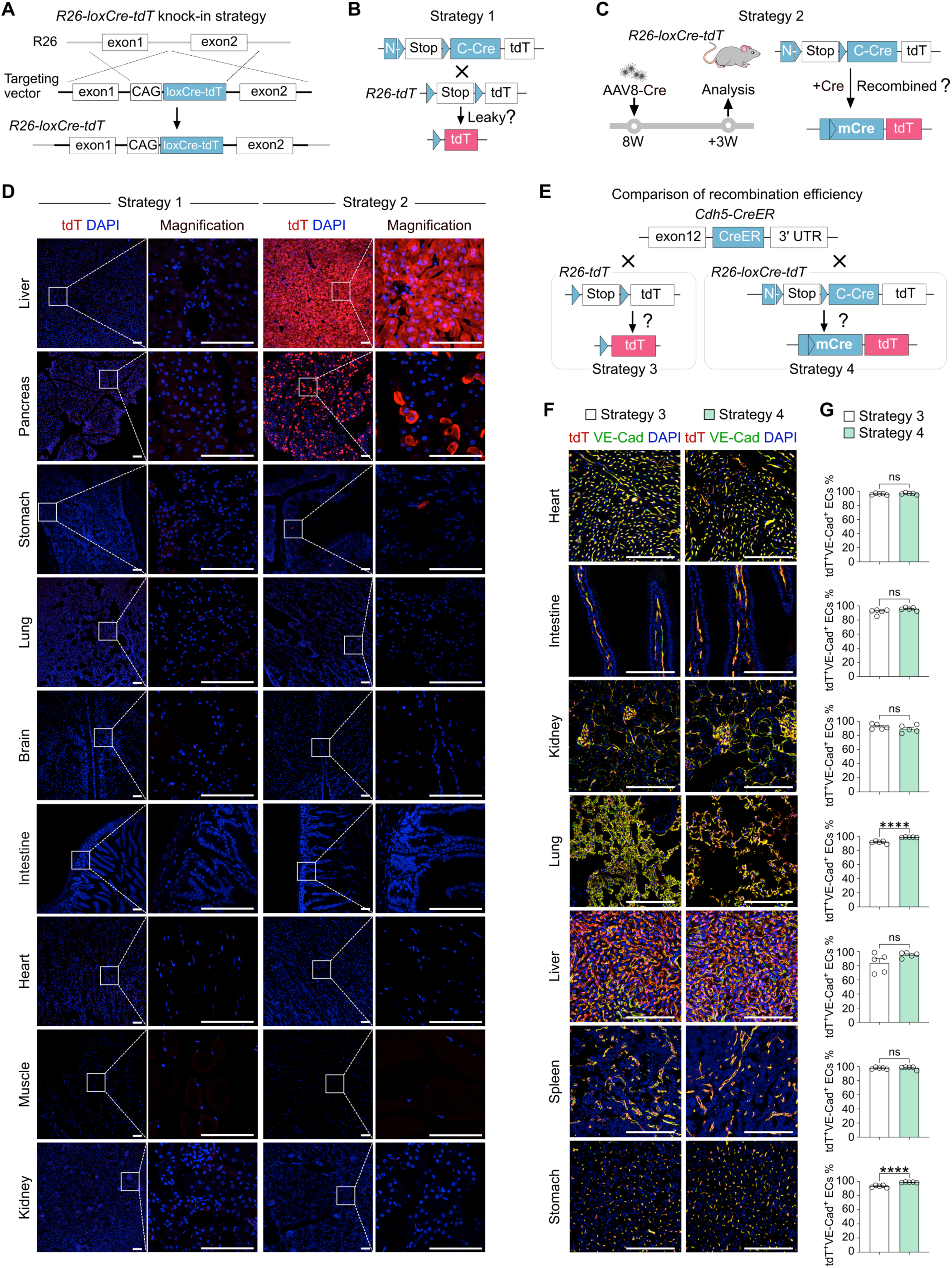
*R26-loxCre-tdT* is efficiently recombined by CreER. (A) A schematic showing the knock-in strategy for the generation of the *R26-loxCre-tdT* allele. In this line, tdT is used to denote the mCre expression after the removal of *Stop*. (B) A schematic showing experimental strategy 1 to examine the leakiness of *R26-loxCre-tdT* mice. (C) A schematic showing experimental strategy 2 to test mCre/tdT expression by AAV8-TBG-Cre. (D) Immunostaining for tdT on tissue sections shows tdT expression in the liver and pancreas when recombination was initiated by AAV8-TBG-Cre. Scale bars, white 100 µm. Each image is representative of 5 individual biological samples. (E) A schematic showing the experimental strategies 3 and 4 for comparing the CreER-mediated first recombination efficiency between *Cdh5-CreER;R26-tdT* and *Cdh5-CreER;R26-loxCre-tdT* mice. (F) Immunostaining for tdT and VE-Cad on liver sections collected on day 7 post-Tam. Scale bars, white 100 µm. Each image is representative of 5 individual biological samples. (G) Quantification of the percentage of VE-Cad^+^ ECs expressing tdT. Data are the means ± SEM; n= 5. \*\*\*\**P* < 0.0001 by student’s *t* test.

We then validate whether the new design can improve the labeling efficiency compared to the conventional approach by crossing endothelial cell-specific *Cdh5-CreER* with the conventional reporter *R26-tdT* (Strategy 3) and the new reporter *R26-loxCre-tdT* mice (strategy 4) under the same protocol of Tam treatment (Figure 4E). Notably, immunostaining and FACS results showed that *R26-loxCre-tdT* significantly limited the leakiness of adult *Cdh5-CreER* compared to *R26-tdT* (Figure S7B). For the labeling efficiency comparison, immunostaining for tdT and VE-Cad on tissue sections revealed no difference in the percentage of VE-Cad^+^ endothelial cells expressing tdT between strategies 3 and 4 (Figure 4F-G), suggesting that recombination of the *R26-loxCre-tdT* allele by CreER was comparable to *R26-tdT*, which is an easy-to-recombine reporter mouse line. Therefore, these data indicate that *R26-loxCre-tdT* can be an easily used reporter, enabling efficient recombination to generate mCre/tdT in CreER-expressing cells without spontaneous leakiness.

### *R26-loxCre-tdT* enables CreER to recombine *R26-Confetti* efficiently

We next examined whether mCre released from loxCre could also display high recombination efficiency for the inert-to-recombine allele *R26-Confetti*. We generated the triple knock-in *Cdh5-CreER;R26-loxCre-tdT;R26-Confetti* mouse line in which mCre was released for confetti recombination upon Tam treatment, and *Cdh5-CreER;R26-Confetti* was used as the control (Figure 5A). Both mouse strains were treated with Tam at 8 weeks of age and their tissue samples were collected for analysis at one week post-Tam (Figure 5B). As tdT expression is also detected in *R26-loxCre-tdT* after the first recombination by CreER, we only compared the YFP and mCFP fluorescence signals of *R26-Confetti* (Figure 5A). Wholemount-fluorescence imaging showed markedly more YFP and mCFP signals in the retina of the *Cdh5-CreER;R26-loxCre-tdT;R26-Confetti* mice, compared to *Cdh5-CreER;R26-Confetti* mice (Figure 5C). Immunostaining on frozen sections revealed a significantly higher percentage of endothelial cells expressing YFP/mCFP in various vascularized organs of the *Cdh5-CreER;R26-loxCre-tdT;R26-Confetti* mice, compared to *Cdh5-CreER;R26-Confetti* mice (Figure 5D). Notably, the results showed that tdT^+^ endothelial cells highly simultaneously expressed YFP/mCFP in *Cdh5-CreER;R26-loxCre-tdT;R26-Confetti* mice (Figure 5D). Thus, these data demonstrate that the *R26-loxCre-tdT* line could be used as an adaptor to enhance CreER-mediated recombination with a simultaneous expression of tdT as an accurate indicator.

**Figure 5.**
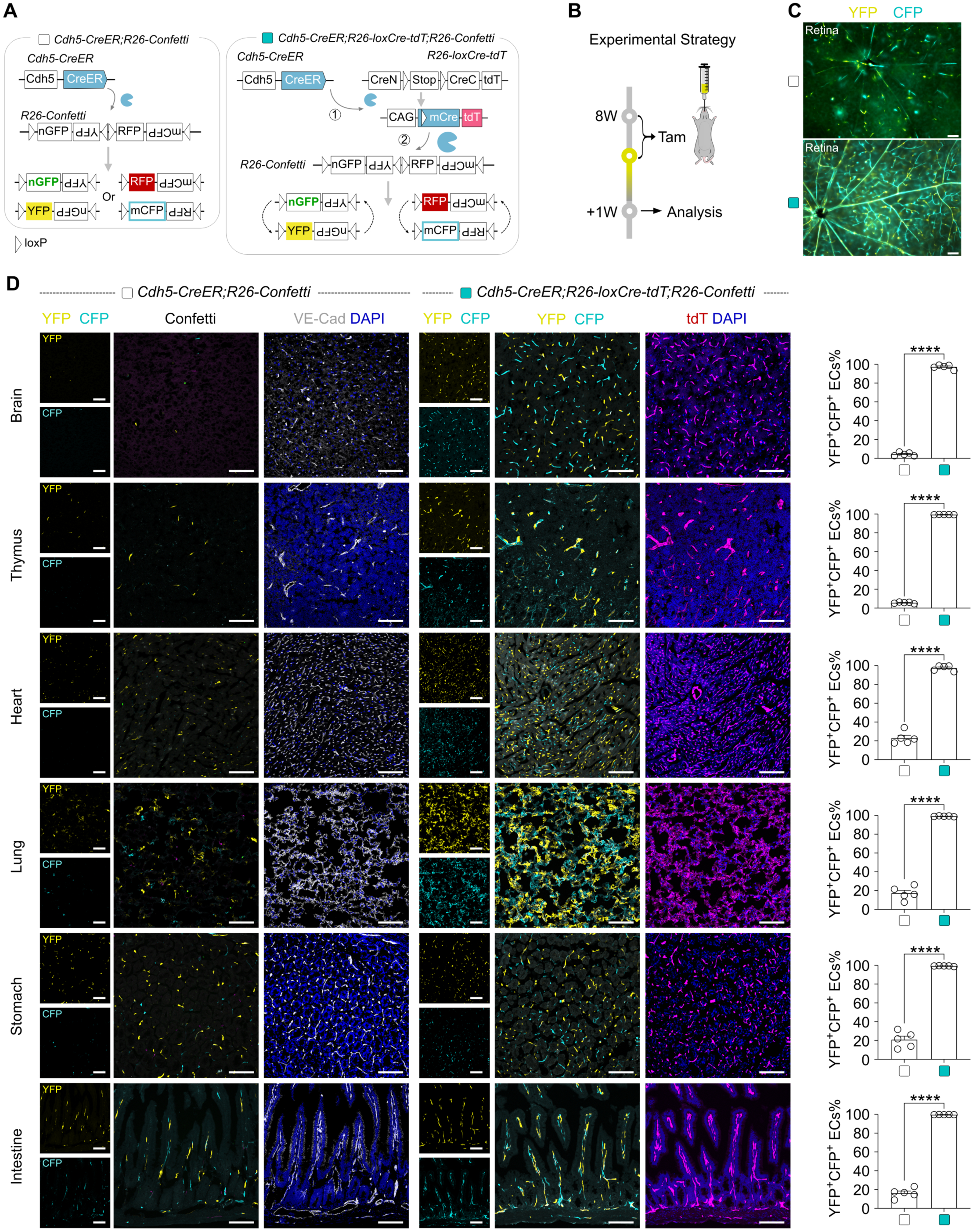
*R26-loxCre-tdT* enables CreER to recombine *R26-Confetti* efficiently. (A) Schematics showing the experimental design. In *Cdh5-CreER;R26-loxCre-tdT;R26-Confetti* mice, Tam-induced CreER-loxp recombination switches *R26-loxCre-tdT* into *R26-mCre-tdT* allele with simultaneous tdT labeling and expression of mCre, which subsequently targets *R26-Confetti* (right panel). The conventional *Cdh5-CreER;R26-Confetti* mice are used as control. (B) A schematic showing the experimental strategy. (C) Whole-mount YFP and mCFP fluorescence images of retina collected from two mice groups. Scale bars, white 100 µm. Each image is representative of 5 individual biological samples. (D) Immunofluorescence images of YFP, mCFP, and VE-Cad on tissue sections show significantly more YFP^+^ and/or mCFP^+^ ECs in the *Cdh5-CreER;R26-loxCre-tdT;R26-Confetti* mice compared with that of *Cdh5-CreER;R26-Confetti* mice (left panel). The right panel shows the quantification of ECs expressing YFP and/or mCFP. Data are the means ± SEM; n = 5. \*\*\*\**P* < 0.0001 by student’s *t* test. Scale bars, white 100 µm. Each image is representative of 5 individual biological samples.

### *R26-loxCre-tdT* adaptor ensures efficient recombination

To evaluate the specificity and efficiency of the loxCre strategy, we generated the *Alb-CreER;R26-loxCre-tdT;R26-Confetti* mouse line, in which *Alb-CreER* recombines *R26-loxCre-tdT* allele in hepatocytes upon Tam treatment to release mCre/tdT and mCre is expected to subsequently recombine the *R26-Confetti* allele (Figure 6A, the right panel). As a control, we crossed the *Alb-CreER;R26-Confetti* mouse with an alternative version of tdT reporter, *R26-tdT2* mice^32^, in which a longe *Stop* cassette was inserted between two loxP sites to reduce the false-positive tracing results (Figure 6A, the left panel). Moreover, we also crossed *Alb-CreER;R26-Confetti* with *iSuRe-Cre* (*R26-loxP-NphiM-Stop-loxP-tdT-Int-Cre*)^18^, in which the first recombination releases tdT and Cre expression, which is expected to subsequently recombine *R26-Confetti* (Figure 6A, the middle panel). The *iSuRe-Cre* is a published and widely utilized inducible reporter-Cre mouse line, which has been reported to improve recombination activity in tdT^+^ cells^18^.

**Figure 6.**
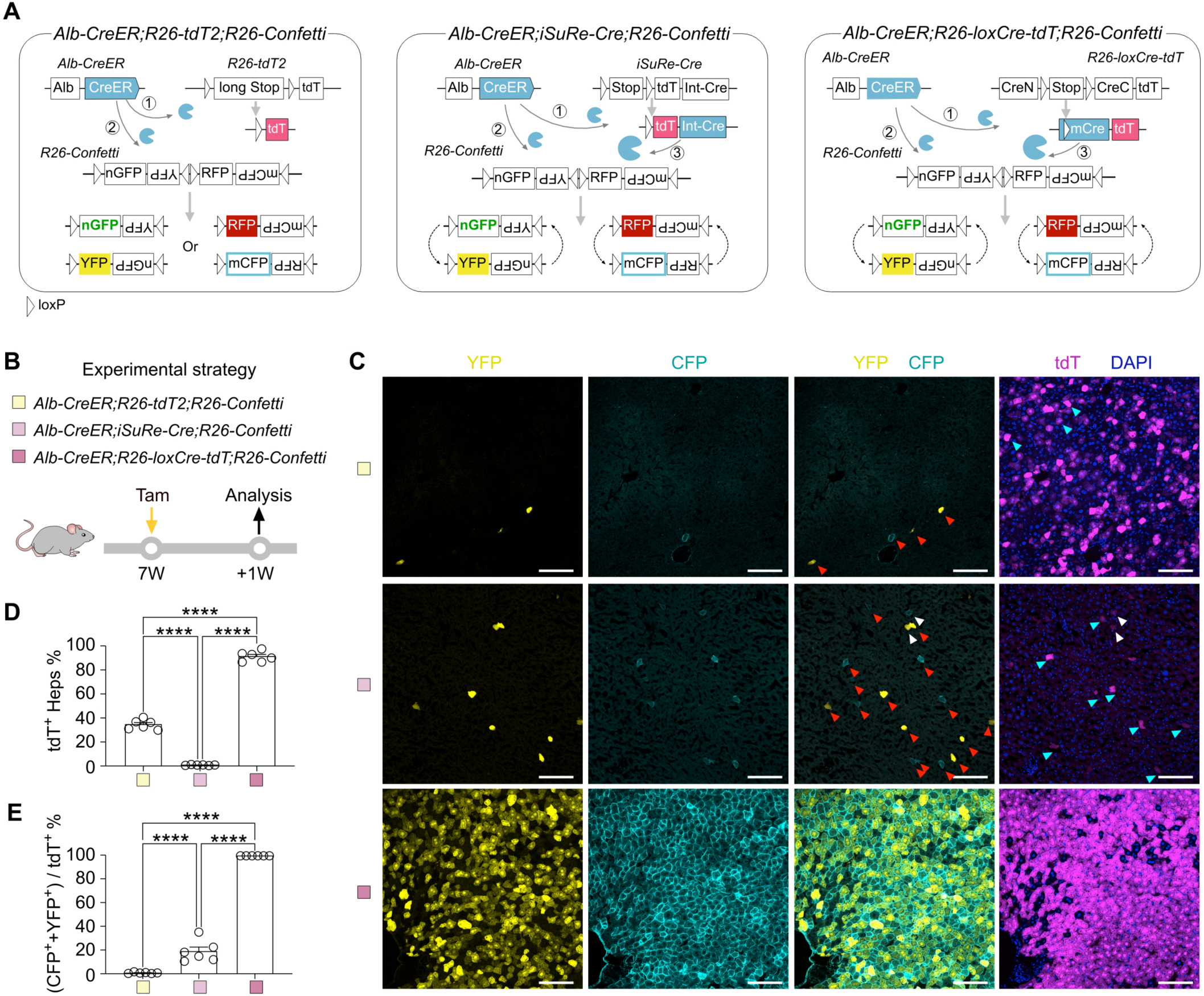
*R26-loxCre-tdT* adaptor ensures efficient recombination. (A) The working strategies for *Alb-CreER;R26-tdT2;R26-Confetti*, *Alb-CreER;iSuRe-Cre;R26-Confetti*, and *Alb-CreER;R26-loxCre-tdT;R26-Confetti*. (B) The experimental strategy. (C) Fluorescence images of YFP, mCFP, and tdT on the liver sections. The red arrows point out some tdT^−^Confetti^+^ false-negative hepatocytes. The white arrows point out some tdT^+^Confetti^+^ true-positive hepatocytes. The cyan arrows point out some tdT^+^Confett^−^ false-positive hepatocytes. (D) Quantification of the tdT^+^ hepatocytes. (E) Quantification of the percentage of tdT^+^ hepatocytes expressing YFP and/or mCFP. Data are means ± SEM; n = 6. \*\*\*\**P* < 0.0001 by one-way ANOVA. Scale bars, white 100 µm. Each image is representative of 6 individual biological samples.

Since most hepatocytes are multinucleated and polyploidy, a single hepatocyte would carry multiple copies of the *R26-Confetti* alleles, yielding two or more fluorescence reporters. Therefore, we used tdT as the readout on *Alb-CreER* mediated recombination of these three following reporters: *R26-tdT2*, *iSuRe-Cre*, and *R26-loxCre-tdT*; and used YFP/mCFP as readouts on *R26-Confetti* (Figure 6C). We injected one dosage of Tam into *Alb-CreER;R26-tdT2;R26-Confetti*, *Alb-CreER;iSuRe-Cre;R26-Confetti*, and *Alb-CreER;R26-loxCre-tdT;R26-Confetti* mice at 7 weeks old and collected their livers for analysis at one week post-Tam (Figure 6B). The fluorescence images presented that the percentage of hepatocytes expressing tdT was highest in *R26-loxCre-tdT* (91.76 ± 1.77%), followed by *R26-tdT2* (35.24 ± 1.66%), and least in *iSuRe-Cre* (1.40 ± 0.13%) (Figure 6D), indicating the orders of recombination efficiency for these alleles were *R26-loxCre-tdT* > *R26-tdT2* > *iSuRe-Cre*. Furthermore, we compared the recombination efficiency on *R26-Confetti* allele and found that the percentage of hepatocytes expressing YFP/mCFP was 1.08 ± 0.23% by *Alb-CreER;R26-tdT2*, 19.42± 3.71% by *Alb-CreER;iSuRe-Cre*; and 100.00 ± 0.00% by *Alb-CreER;R26-loxCre-tdT* (Figure 6E). Together, these findings collectively indicate that the mCre, when released from the *R26-loxCre-tdT* construct, exhibits a significantly enhanced recombination efficiency. This improvement is notable when compared to both the conventional CreER and previously reported *iSuRe-Cre*^18^.

### *R26-loxCre-tdT* ensures gene deletion in tdT^+^ cells

To investigate whether *R26-loxCre-tdT* enables *Alb-CreER* to more efficiently delete genes, we crossed *Alb-CreER;R26-loxCre-tdT* with *Ctnnb1^flox/flox^* to generate *Alb-CreER;R26-loxCre-tdT*;*Ctnnb1^flox/flox^* mice for evaluating the efficiency of releasing mCre and also mCre-mediated *Ctnnb1* deletion in tdT^+^ hepatocytes (Figure 7A). We also crossed *Alb-CreER;R26-tdT2* with *Ctnnb1^flox/flox^* to generate *Alb-CreER;R26-tdT2;Ctnnb1^flox/flox^* mice to evaluate recombination efficiency of both *R26-tdT2* and *Ctnnb1^flox/flox^*alleles (Figure 7A). Additionally, *Alb-CreER;R26-loxCre-tdT*;*Ctnnb1^flox/+^* mice were used as a control for heterozygous gene deletion (Figure 7B). Firstly, the FACS and immunostaining data showed no leakiness in *Alb-CreER;R26-loxCre-tdT* (Figure S8A). Based on the no leaky background, we treated *Alb-CreER;R26-tdT2;Ctnnb1^flox/flox^* and *Alb-CreER;R26-loxCre-tdT*;*Ctnnb1^flox/+^* mice with five dosages of Tam, and treated *Alb-CreER;R26-loxCre-tdT;Ctnnb1^flox/flox^*mice only with one dosage of Tam, and collected hepatocytes for analyzing *Ctnnb1* gene deletion at 4 weeks after Tam treatment (Figure 7B). Immunostaining data showed that β-catenin, GS, and E-CAD were significantly reduced in *Alb-CreER;R26-loxCre-tdT;Ctnnb1^flox/flox^*mice but were readily detectable in both *Alb-CreER;R26-tdT2;Ctnnb1^flox/flox^*and *Alb-CreER;R26-loxCre-tdT*;*Ctnnb1^flox/+^* mice (Figure 7C). Western blotting of β-catenin and qRT-PCR of *Ctnnb1* of isolated hepatocytes revealed the significant deletion of *Ctnnb1* in the *Alb-CreER;R26-loxCre-tdT;Ctnnb1^flox/flox^*mice compared with those in control groups (Figure 7D-F). Additionally, Wnt downstream or regulated genes, including *Glul*, *Axin2*, *Cyp2e1*, *Oat*, *Tcf7*, *Lect2*, *Tbx3*, and *Slc1a2* were significantly reduced in hepatocytes derived from the *Alb-CreER;R26-loxCre-tdT;Ctnnb1^flox/flox^*mice, compared with those from the *Alb-CreER;R26-tdT2;Ctnnb1^flox/flox^* and *Alb-CreER;R26-loxCre-tdT*;*Ctnnb1^flox/+^* mice, respectively (Figure 7G).

**Figure 7.**
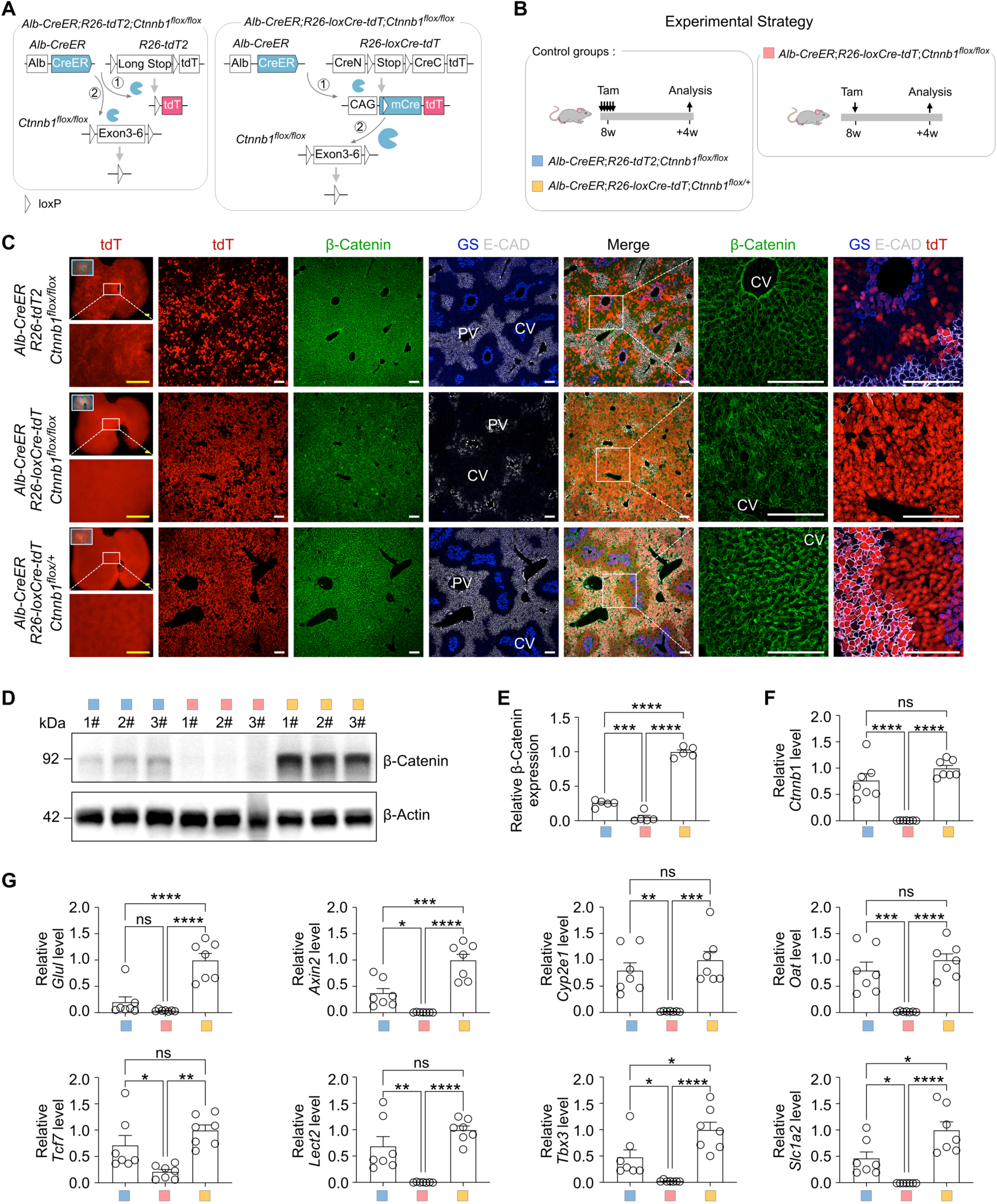
*R26-loxCre-tdT* enables *Alb-CreER* to efficiently knockout genes in hepatocytes. (A) Schematics showing the experimental designs for *Ctnnb1* gene knockout using either *Alb-CreER;R26-tdT2* or *Alb-CreER;R26-loxCre-tdT* mice. (B) A schematic showing the experimental strategy. *Alb-CreER;R26-tdT2;Ctnnb1^flox/flox^* and *Alb-CreER;R26-loxCre-tdT;Ctnnb1^flox/+^* mice were injected with Tam for 5 times, while *Alb-CreER;R26-loxCre-tdT;Ctnnb1^flox/flox^* mice were injected with Tam for one time. (C) Immunostaining for tdT, β-Catenin, GS, and E-CAD on liver sections. Scale bars, yellow 1mm; white 100 µm. Each image is representative of 5 individual biological samples. (D) Western blotting of β-Catenin and β-Actin expression in hepatocytes, n = 3. (E) Quantification of β-Catenin expression. Data are the means ± SEM; n = 5. (F) qRT-PCR analysis of the relative expression of *Ctnnb1* in the hepatocytes. (G) qRT-PCR analysis of the relative expression of *Glul*, *Axin2*, *Cyp2e1*, *Oat*, *Tcf7*, *Lect2*, *Tbx3*, and *Slc1a2* in hepatocytes. Data are the means ± SEM; n = 7; ns, non-significant; \**P* < 0.05, \*\**P* < 0.01, \*\*\**P* < 0.001, \*\*\*\**P* < 0.0001 by one-way ANOVA.

We next elucidate whether the significant superiority of *Alb-CreER;R26-loxCre-tdT;Ctnnb1^flox/flox^* in the above data is attributed to low Tam dosage, which may reduce Tam toxicity and side effects. Initially, we treated *Alb-CreER;R26-tdT;Ctnnb1^flox/flox^*mice with one dosage of Tam and subsequently sorted out tdT^+^ hepatocytes for qRT-PCR against *Ctnnb1* 4 weeks after Tam treatment. The results indicated that *Alb-CreER* with a single dose of Tam, without the assistance of *R26-loxCre-tdT*, was insufficient to knock out *Ctnnb1* (Figure S8B-D). Next, we conducted an experiment involving a proportional gradient of the Tam dosages, in which we gave five dosages, one dosage, one-fifth of one dosage, one-twenty-fifth of one dosage, and one-hundred-twenty-fifth of one dosage of Tam to *Alb-CreER;R26-loxCre-tdT;Ctnnb1^flox/flox^*mice, respectively. We analyzed samples 3 weeks after Tam’s treatment. The FACS results showed that five groups of the proportional gradient of the Tam dosages labeled a gradient of tdT^+^ cells: 94.33±0.76%, 90.14±0.41%, 57.07±0.51%, 27.30±2.35%, and 1.27±0.54% (Figure S8E), respectively. We sorted out tdT^+^ hepatocytes from each group for qRT-PCR against *Ctnnb1*. Our finding revealed that varying doses of Tam treatment primarily affect the target cell ratio but do not influence the high recombination efficiency of mCre at loxP sites within the *Ctnnb1* allele (Figure S8E-F). Additionally, the immunostaining analysis revealed a noticeable trend, in which the proportion of GS^+^ or E-CAD^+^ hepatocytes was in decreased tendency with the increase of target tdT^+^ hepatocytes (Figure S8F-G). This observation can be attributed to the significant knockout of *Ctnnb1* in tdT^+^ cells (Figure S8E-F).

Together, the combined results lead to the conclusion that *R26-loxCre-tdT* significantly enhances the recombination efficiency of CreER, facilitating the targeting of genes that are challenging to manipulate by traditional CreER-loxP recombination. Furthermore, the tdT component in *R26-loxCre-tdT* enables precise tracing of the modified cell lineage, which accurately investigates the roles of genes in various biological processes.

## Discussion

In this study, we demonstrated that the *rox*-*Stop-rox* insert placed in front of *Cre* does not effectively prevent the leakiness of Cre in the Rosa26 allele. To address this issue, we modified *Cre* by inserting the *RSR* cassette into its coding region, thereby dividing Cre into its N- and C-terminal segments. Our study showed that the developed roxCre not only eliminates leakage but also enables the release of mCre for targeted gene manipulation in specific cell subpopulations through the promotion of DreER. According to the construction of roxCre, we advanced to develop *R26-loxCre-tdT*, which significantly enhances the efficiency of inducible Cre-mediated gene manipulation, particularly when employing less effective CreER drivers for gene functional exploration.

Assuming successful gene deletion in the cell type of interest depends on efficient cell labeling by a lineage-specific reporter mediated by the same Cre is unrigorous, as recombination efficiency varies significantly among different mouse lines. Our data clearly showed the differences in recombination efficiency of different alleles (e.g. *R26-tdT*, *R26-tdT2*, and *R26-Confetti*), even located in the same genomic locus (e.g. Rosa26) mediated by the same CreER under the same Tam treatment protocol (Figure 6, Figure 7C, and Figure S8D). However, *R26-loxCre-tdT* can be efficient for recombination mediated by CreER and release mCre in targeted cells, thus enabling efficient and accurate manipulating of cells of interest.

Compared to the previously reported transgenic mice line, *iSuRe-Cre*^18^, *R26-loxCre-tdT* shows more stable and robust enhancement in the efficiency of CreER recombination (Figure 6, Figure 7), avoiding the toxic effects resulting from high-dose of Tam. This advancement significantly improves the reliability of genetic manipulation and cell fate lineage tracing, making *R26-loxCre-tdT* a valuable tool for researchers investigating cellular functions and behaviors. The research group of the iSuRe-Cre system recently reported a new system, named iSuRe-HadCre, which demonstrated to be effective in deleting floxed genes while avoiding the toxicity of constitutive Cre^33^ Further studies can be carried out to compare the recombination enhancement between *R26-loxCre-tdT* and *iSuRe-HadCre*. Given the high recombination efficiency of mCre, we envision that loxCre can be used for mosaic analysis, where multiple alleles in individual cells need to be efficiently recombined for evaluation of cell-autonomous gene function with genetic labeling.

Importantly, Cre toxicity represents a substantial concern. The toxicity may stem from the administration of Tam/4-OHT and the constitutive expression of Cre driven by specific promoters, including *CD4*, *Ins2*, and *Myh6*^15^. Cre toxicity has been shown to impact diverse cellular processes, including DNA damage, cell proliferation, cell apoptosis, inflammation, metabolic signaling, and genetic dysfunction^14, 15^. Although roxCre and loxCre can mitigate the toxicity associated with high-dose Tam, they cannot eliminate the Cre toxicity, as mCre is constitutively expressed following recombination. Considering the lack of Cre toxicity evaluation of the newly developed mouse lines in this study, Cre toxicity should be taken into account in future biological functional studies involving roxCre and loxCre systems. Furthermore, to optimize the robustness and rapidity of Cre expression while minimizing the toxic effects associated with constitutive Cre, further development of iterated Cre removal system should be carried out for both loxCre and roxCre systems in the future.

Wnt-β-catenin plays an important role in liver development, zonation, homeostasis, and diseases^30^. The administration of five doses of Tam in *Alb-CreER* mice did not impact downstream protein GS expression due to not knocking out *Ctnnb1* completely. However, a single dose of Tam could effectively induce a significant knockout of *Ctnnb1* through *Alb-CreER;R26-loxCre-tdT* (Figure 7). This indicates that *R26-loxCre-tdT* can assist CreER in addressing challenging recombined loxP sits. Notably, *R26-loxCre-tdT* can release mCre by CreER-mediated activation and report tdT expression across all cells that express the recombinase. Therefore, this system does not produce a gradient of reporter fluorescence reflective of the promoter activity of CreER. Consequently, it’s advisable not to use *R26-loxCre-tdT* with a nonspecific and unstable CreER mouse line, which would amplify the CreER mouse line’s weakness and result in unanticipated nonspecific cell-targeting.

Furthermore, the strategy of roxCre further enhances the precision for cell fate mapping with genetic deletion simultaneously in Dre^+^Cre^+^ cells through intersectional genetic targeting. For example, we have showcased the strength of roxCre for genetic manipulation in peri-central hepatocytes, resulting in efficient gene knockout in tdT^+^ hepatocytes specifically (Figure 3). Additional roxCre mouse lines driven by different promoters can serve as valuable tools for detailed investigations into specific cell subpopulations. By using roxCre, studies can achieve precise spatial and temporal control of gene expression, facilitating targeted analysis that contributes to the knowledge of developmental biology, disease mechanisms, and potential therapeutic interventions.

Previous studies have used Cre-loxP to simultaneously over-express gene and fluorescence reporter in a cell for functional genetic mosaic (ifgMosaic) analysis^34^. However, ifgMosaic could not take advantage of the abundant resources from existing available floxed mouse lines for loss-of-function study. Mosaic mutant analysis with spatial and temporal control of recombination (MASTR) utilized FlpoER for tissue-specific expression of GFPCre to delete floxed genes^16^. Compared with FlpoER lines, CreER is a more broadly used recombinase that most laboratories use in gene deletion experiments. A recent study using dual recombinase-mediated cassette exchange (MADR) also permits stable labeling of mutant cells expressing transgenic elements from defined chromosomal loci^35^. However, the use of viruses and electroporation, albeit convenient and rapid, are inefficient to specifically target any type of cells for *in vivo* studies. The powerful mosaic analysis with double markers (MADM) enables simultaneous lineage tracing of a pair of mutant and control sibling cells with distinct fluorescence reporters, allowing precise mosaic analysis of gene function in any cell^36, 37^. Nevertheless, the MADM cassette has to be combined with the mutant null allele, which is not readily available for synchronizing reporter expression and genetic modification in cells of interest. The ensured gene deletion with tdT reporter by *R26-loxCre-tdT* could be coupled with present CreER and loxP alleles for potential mosaic analysis. This could be achieved by adjusting Tam at a lower dosage so that the recombination efficiency to release mCre could be low in cells of interest for sparse tdT labeling. We believe that *R26-loxCre-tdT* could be useful for robust, efficient, and temporal gene deletion at a population level and mosaic analysis at a single-cell level with improvement in the future.

In conclusion, we developed two novel genetic tools, roxCre and loxCre, permitting efficient recombination to release mCre, which greatly enhanced the effectiveness of subsequent gene deletion in tdT^+^ cells, thus facilitating further detailed characterization of these cells both *in situ* and *ex vivo*. These tools are particularly beneficial when utilizing less effective DreER or CreER drivers to better elucidate the roles of specific genes in biological processes, facilitating more robust studies.

## Materials and Methods

### Mice

Experiments using mice (Mus musculus) were carried out with the study protocols (SIBCB-S374-1702-001-C1) approved by the Institutional Animal Care and Use Committee of Center for Excellence in Molecular Cell Science (CEMCS), Shanghai Institute of Biochemistry and Cell Biology, Chinese Academy of Sciences. The *R26-DreER*^38^, *Alb-CreER*^26^, *R26-GFP*^29^, *R26-tdT*^11^, *R26-tdT2*^32^, *R26-RSR-tdT*^39^, *R26-Confetti*^12^, *iSuRe-Cre*^18^ and *Ctnnb1-flox*^31^ mouse lines were used as previously described. New knock-in mouse lines *R26-RSR-Cre*, *R26-RSR-Cre2*, *R26-R-reverseCre-R*, *Cdh5-CreER*, *Alb-roxCre1-tdT*, *Alb-roxCre7-GFP*, *Cdh5-roxCre4-tdT*, *Cdh5-roxCre10-GFP*, *Cyp2e1-DreER*, and *R26-loxCre-tdT* were generated by homologous recombination using CRISPR/Cas9 technology. These new mouse lines were generated by the Shanghai Model Organisms Center, Inc. (SMOC). These mice were bred in a C57BL6/ICR mixed background. All mice were housed at the laboratory Animal center of the Center for Excellence in Molecular Cell Science in a Specific Pathogen Free facility with individually ventilated cages. The room has controlled temperature (20–25 °C), humidity (30–70%), and light (12 hours light-dark cycle). For the determination of the embryonic period of sampling, the day on which the vaginal plug was examined in female mice was recorded as E0.5. As the sex is not relevant to the topic of this study, male and female mice ranging in age from E13.5 to 12W (week) were allocated and mixed into the experimental groups in this study. No data in mouse experiments were excluded.

### Genomic PCR

Genomic DNA was prepared from the mouse toes or embryonic tails. Tissues were precipitated by centrifugation at maximum speed for 1 min at room temperature. After that, tissues were lysed by lysis buffer (100 mM Tris-HCl, 5 mM EDTA, 0.2% SDS, 200 mM NaCl, and 100 μg/mL Proteinase K) at 55 °C overnight. About 750μL pure ethanol was added to the lysis mixture and mixed thoroughly, followed by centrifugation at maximum speed for 5 min at room temperature to collect the DNA precipitation. Then the supernatant was discarded and the mixture was dried at 55 °C for 1 hour. About 250μL double-distilled H_2_O was added to dissolve the DNA. The genomic PCR primer pairs were designed for the mutant alleles spanning both endogenous genomic fragments and insert fragments. The genetic constructs and genotyping of the new knock-in lines can be reviewed in Figure S9.

### *In vitro* screening of mCre for efficient recombination

To test if the remaining rox sequence after Dre-rox recombination would impact Cre activity, the pCDNA3.1^40^ (Invitrogen, V79020) was used to express 12 types of modified Cre (mCre). For testing the Cre activity, 500ng pCDNA3.1-mCre plasmids, or pCDNA3.1 (negative control), or pHR-CMV-nlsCRE (positive control, Addgene, 12265) were mixed with 500ng pCAG-loxp-stop-loxp-ZsGreen (Addgene, 51269) plasmid. *In-vitro* transfection experiment protocol is performed according to that described previously^41^. For cell culture medium, DMEM (ThermoFisher, 11965092) was supplemented with 10% fetal bovine serum (Gibco, 10099141c) for preparing fresh complete culture medium. Poly-D-Lysine (ThermoFisher, A3890401) precoated coverslips (Biosharp, BS-14-RC) or 10 cm dish (ThermoFisher, 150466) overnight to dry, and sterile water was used to flush it for three times. A cryogenic vial of QBI293 was put in a 37°C water bath, then thawed liquid contents into a 50 mL conical tube (Corning,430829) prefilled with 5 mL prewarmed fresh complete culture medium. After centrifuging at 125 × g for 5 min, the supernatant was discarded and cells were resuspended in 10 mL complete medium. One-third of the cells were plated in a 10 cm coated dish and incubated in cultures at 37°C. On the second day, the culture medium was discarded, and the cell layer was briefly rinsed with prewarmed PBS (Gibco, C10010500BT). After that, PBS was removed and 1 mL 0.25% Trypsin solution (with EDTA, Gibco, 25200072) was added. The cells were incubated at 37°C for 3 min., and a 6mL complete growth medium was added to aspirate cells by gently pipetting. The medium-containing cells were transferred to a 50 mL conical tube and centrifuged at 125 × g for 5 min. The supernatant was discarded, and cells were resuspended in 5.2 mL complete culture medium. 24 well plates were placed into coated coverslips and added 1mL complete culture medium. 200µL cells were added to every experimental well and incubated cultures at 37°C for 9 hours to grow up to ∼80%. The new prewarmed fresh complete culture medium replaced the old for 1mL per well. Lipofectamine™ 3000 Transfection Reagent (Thermo Fisher, L3000015) was used for plasmid transfection. Samples were incubated in cultures at 37°C for 24 hours. Samples were washed by PBS once and mounted on slides by VECTASHIELD^®.^ Antifade Mounting Medium with DAPI (Vector, H-1200). Images were acquired using an Olympus microscope (Olympus, BX53).

### Tamoxifen injection and sample collection

For induction of CreER or DreER, tamoxifen (Sigma, T5648) was dissolved in corn oil and was administered to mice by oral gavage. *Cyp2e1-DreER;Alb-roxCre1-tdT;Ctnnb1^flox/+^*and *Cyp2e1-DreER;Alb-roxCre1-tdT;Ctnnb1^flox/flox^* mice were treated with 0.05 mg Tam per gram mouse body weight (0.05 mg/G) for one dose. For comparing tdT^+^ cells, *Cyp2e1-DreER;R26-RSR-tdT* and *Cyp2e1-DreER;Alb-roxCre1-tdT* mice were treated with 0.05 mg/G Tam. *R26-DreER;Alb-roxCre1-tdT;R26-Confetti*, *R26-DreER;Cdh5-roxCre4-tdT;R26-Confetti*, *R26-DreER;Alb-roxCre7-GFP;R26-Confetti*, *R26-DreER;Cdh5-roxCre10-GFP;R26-Confetti*, *R26-DreER; R26-DreER;Alb-roxCre7-GFP;R26-GFP*, *Alb-CreER;R26-Confetti*, *Cdh5-CreER;R26-Confetti*, *Alb-CreER;R26-tdT2;R26-Confetti*, *Alb-CreER;iSuRe-Cre;R26-Confetti*, *Alb-CreER;R26-loxCre-tdT;R26-Confetti*, *Alb-CreER;R26-loxCre-tdT;Ctnnb1^flox/flox^*, and *Alb-CreER;R26-tdT;Ctnnb1^flox/flox^* mice were treated with 0.2 mg/G Tam for one dose. *Cdh5-CreER;R26-tdT*, *Cdh5-CreER;R26-loxCre-tdT*, *Cdh5-CreER;R26-Confetti*, *Cdh5-CreER;R26-loxCre-tdT;R26-Confetti*, *Alb-CreER;R26-tdT2;Ctnnb1^flox/flox^*, and *Alb-CreER;R26-loxCre-tdT;Ctnnb1^flox/+^*mice were treated with 0.2 mg/G Tam one dose per day for 5 days. *R26-RSR-Cre2;R26-GFP* and *R26-R-reverseCre-R;R26-tdT* mice were sacrificed at 8 weeks old for analysis. For the proportional gradient of the Tam dosages, five dosages group was given 0.2 mg/G Tam for constitutive 5 days, one-fifth of one dosage group was given 0.04 mg/G Tam for one dose, one-twenty-fifth of one dosage group was given 0.008 mg/G Tam for one dose, and one-hundred-twenty-fifth of one dosage group was given 0.0016 mg/G Tam for one dose. Mice, both males and females, at the age of 8–12 weeks were used for experiments with similar-aged mice for both control and experimental groups.

### Whole-mount imaging and sectioning

The tissue samples were fixed with 4% paraformaldehyde (PFA) (Sigma, P6148-500g, wt/vol in PBS) for 1 or 2 hours (stomach, intestine, colon) at 4°C, followed by washing with PBS three times. The fixed tissues were placed in an agarose-filled petri dish for bright-field and fluorescence imaging by a Zeiss stereoscopic microscope (AxioZoom V16). For cryo-sections, tissues were sectioned to slides of 10-μm thickness after dehydration by 30% sucrose (Sinopharm, H-10021463, wt/vol in PBS) overnight and pre-embedding with OCT (Scigen, 4586) at 4 °C for 1 hour.

### Immunostaining

Immunostaining was performed as previously described^42^. 0.2% PBST was prepared with 0.2% (vol/vol) Triton X-100(Sigma, T9284) dissolved in PBS. Tissue sections were blocked with 2.5% normal donkey serum and 0.1% 4’6-diamidino-2-phenylindole (DAPI, Invitrogen, D21490) dissolved in 0.2% PBST for 30 mins after washing with PBS three times. The tissue sections were incubated with primary antibody diluted in 0.2% PBST at 4°C overnight. On the next day, sections were incubated with secondary antibodies diluted in 0.2% PBST at room temperature for 30 min, followed by PBS washing for three times. The slides were washed with PBS for three times. The slides were mounted with a mounting medium (Vector Lab). For weak signals, the endogenous peroxidase activity was quenched before blocking. Horseradish peroxidase or biotin-conjugated secondary antibodies and a Tyramide signal amplification kit (PerkinElmer) were used after incubating the primary antibodies. For primary antibodies of murine origin, mouse immunoglobulins were blocked with an anti-mouse Fab antibody (Jackson, 715-007-003, 1:100). The included primary antibodies are listed as follows: tdT (Rockland, 600-401-379, 1:500; or Rockland, 200-101-379, 1:500), GFP (Invitrogen, A11122, 1:500), GFP (Rockland, 600-101-215M, 1:500), GFP (Nacalai, 04404-84; 1:500), GS (Abcam, ab49873, 1:1000), β-catenin (BD Pharmingen, 610153, 1:200), E-CAD (R&D, AF748, 1:500), Ep-CAM (Abcam, ab92382, 1:500), VE-Cad (R&D, AF1002, 1:100), and CYP2E1(Abcam, ab28146, 1:100). The corresponding secondary antibodies (JIR or Abcam) were diluted according to the instructions. Images were captured by using a Nikon confocal (Nikon A1 FLIM) or an Olympus confocal (FV3000), and captured images were analyzed by Image J2 (version 2.9.0/1.54f) and Photoline (version 23.02) software. We collected five random fields from each liver section for quantification. Mutant and control sections were processed at the same time to avoid potential batch differences during staining. Imaging of all immunostained slides was performed under the same exposure and contrast conditions using the same confocal microscope.

### Imaging of Confetti fluorescence reporters

All the fluorescence images of *R26-confetti* were taken by a Leica confocal (Leica SP8 will). All the sections were collected freshly and blocked with 0.2% PBST containing DAPI for 30 mins after washing with PBS three times. The parameters of excitement light and emission light for all channels are set as follows: DAPI (ex.405nm, em.415-450nm), CFP (ex.458nm, em.463-481nm), GFP (ex.488nm, em.495-508nm), YFP (ex.514nm, em.520-535nm), tdT (ex.546nm, em.555-590nm), markers for antibody staining (ex.647nm, em.657-700nm). Captured images were analyzed by Image J2 (version 2.9.0/1.54f) and Photoline (version 23.02) software.

### Hepatocyte dissociation

Mouse primary hepatocytes were isolated by the two-step collagenase perfusion method, which was modified from a previous protocol^43^. Briefly, mice were anesthetized and the liver was exposed through an incision in the lower abdomen. The inferior vena cava and portal vein were also exposed. A needle was inserted into the inferior vena cava and secured with a hemostatic clamp around the needle. The portal vein was cut immediately when the mouse liver was perfused with a perfusion medium buffer (containing 0.5 mM EGTA) for 5 minutes using a peristaltic pump. Then, the liver was perfused with medium containing collagenase type I (150 U/ml; Gibco, 17100-017) for 2-5 min to adequately digest the liver. After the gallbladder was removed, the liver was dissected with cold DMEM to free the hepatic cells. Then the cell suspension was passed through a 70-μm cell strainer (BD Biosciences, 352350) and centrifuged at 50 x g for 3 min at 4 °C. The supernatant was removed, and cells were resuspended in Percoll (GE Healthcare, 17-0891-01)/DMEM/10 x PBS (Sangon, B548117, diluted in 1:1) (1:1:0.1) mixture and centrifuged at 300 x g for 5 min at 4 °C. After the supernatant was removed, cells were dissected with cold DMEM and centrifuged at 50 x g for 3 min at 4 °C. Purified hepatocytes were collected for FACS analysis, qRT-PCR or Western Blot analysis.

### Flow cytometric analysis and isolation of tdT^+^ hepatocytes

Cells were centrifuged at 50 x g for 3 min at 4 °C and dissected with relevant suspended solution. The suspended solution is 0.1% Dnase I (Worthington, LS002139, diluted by DMEM) mixed with 0.1% DAPI. tdT^+^ hepatocytes were sorted by FACS Aria SORP machine and Sony MA900. Hepatocytes were collected by DMEM and then centrifuged at 50 x g for 3 min at 4 °C.

### Hepatic Non-parenchymal cell dissociation

The liver was diced into 1mm pieces and then transferred into 10 mL digestion buffer (0.05g/100 mL collagenase type IV (Worthington, LS004188), 2mg/mL collagenase type I, 5u/mL Dispase (Corning, 354235), and 1%Dnase I in PBS). The sample was incubated in a 37°C incubator shaker at 220 rotations per minute for 30 min, with mixing performed three times during the incubation. Following this, 0.5 mL FBS was added to the digestion mixture. The resulting cell suspension was passed through a 40-μm cell strainer (BD Biosciences, 352340), and transferred to a new t ube containing 30 mL cold PBS after being centrifuged at 50 x g for 3 min at 4 °C. The supernatant was discarded after 700 x g for 5 min at 4 °C. To lyse erythrocytes, RBC lysis buffer was added, and the mixture was kept for at room temperature 5 min. The lysis was halted by adding 20 m L cold PBS, followed by another centrifugation at 700 x g for 5 min at 4 °C. The supernatant was discarded, and the pellet was resuspended in the suspended solution.

### Total RNA extraction and qRT-PCR

Total RNA was extracted from hepatocytes isolated from indicated mice as previously described^44^. Cells were lysed with Trizol (Invitrogen, 15596018), and total RNA was extracted according to the manufacturer’s instructions. For each sample, 1 μg of total RNA was reversely transcribed into cDNA using the Prime Script RT kit (Takara, RR047A). The SYBR Green qPCR master mix (Thermo Fisher Scientific, 4367659) was used and quantitative RT-PCR was performed on QuantStudio 6 Real-Time PCR System (Thermo Fisher Scientific). *Gapdh* was used as the internal control. Sequences of all primers would be provided upon request.

### Western blot

All samples were lysed in RIPA lysis buffer (Beyotime, P0013B) containing protease inhibitors (Roche, 11836153001) for 30 min on ice, and then centrifuged at 15,000 x g for 15 min to collect the supernatant. All samples were separated with 30μL for testing the protein concentration by Pierce^TM^ BCA Protein Assay kits (Thermo Scientific^TM^,23227). The remaining samples were mixed with 5 x loading buffer (Beyotime, p0015L) and boiled at 100 °C for 5 min. 40μg samples were added to every well in precast gradient gels (Beyotime, P0469M) with 1x running buffer (Epizyme, PS105S, should be diluted into 1x). After running, samples were transferred onto Immobilon® PVDF membranes (Millipore, IPVH00010). After blocking in blocking buffer (Epizyme, PS108P), the membranes were incubated with primary antibodies (diluted by primary antibody dilution buffer (Epzyme, PS114)) overnight at 4 °C, then washed for three times and incubated with HRP-conjugated secondary antibodies (diluted by 1xTBST (Epzyme, PS103S, should be diluted into 1x)). Samples were incubated with chemiluminescent HRP substrate (Millipore, WBKLS0500), and related signals were detected by MiniChemi 610 Plus (Biogp). The following antibodies were used: β-catenin (BD Biosciences, 610153, 1: 5000), GAPDH (Proteintech, 60004-1-IG, 1:2000), β-actin (Epzyme, LF202,1:1000), HRP-donkey-a-mouse (JIR, 715-035-150, 1:5000), and HRP-goat a rabbit IgG (JIR, 111-035-047, 1:5000).

### AAV injection

The AAVs used in this study were purchased from Taitool Biotechnologies company (Shanghai, China). Briefly, AAVs were produced with cis-plasmids containing the full TBG promoter, which is specifically active in hepatocytes, and Cre expression is under the control of TBG promoter, replication incompetent AAV2/8-TBG-Cre virus (AAV8-TBG-Cre, S0657-8-H5) was packaged and purified before application to mice. Mice were injected intraperitoneally at 1× 10^11^ genome copies of the virus per mouse. The above newly generated virus as well as targeting plasmids will be provided upon request.

### Statistical analysis

Each pot in every graph represented one individual mouse. The quantification of each mouse based on the fluorescence images was performed by counting the average data of five 10x fields from different sections. Statistical analyses were performed using GraphPad Prism (Version 9.5.1). All the continuous variables were expressed as means ± standard error of the mean (SEM). One-way ANOVA was used to detect statistical significance between three experimental groups. The statistical difference between the two experimental groups was determined using the unpaired student’s *t* test. The probability (*P*) values of less than 0.05 were considered statistically significant. The equation, Log(agonist) vs. response – Variable slope (four parameters) in Prism, was adapted for the dose-response curve for the proportional gradient of the Tam dosages experiment analysis, with a confidence interval of 95%.

## Acknowledgments

This study was supported by the National Key Research & Development Program of China (2024YFA1803302, 2023YFA1800700, 2022YFA1104200, 2023YFA1801300, 2020YFA0803202), National Natural Science Foundation of China (82088101, 32370897, 32100648, 32370783, 32100592), CAS Project for Young Scientists in Basic Research (YSBR-012), Shanghai Pilot Program for Basic Research-CAS, Shanghai Branch (JCYJ-SHFY-2021-0), Research Funds of Hangzhou Institute for Advanced Study (2022ZZ01015, B04006C01600515), Shanghai Municipal Science and Technology Major Project, and the New Cornerstone Science Foundation through the New Cornerstone Investigator Program and the XPLORER PRIZE, and CAS-Croucher Funding Scheme for Joint Laboratories.

## Author contributions

M.S., J.L., and B.Z. designed the study, analyzed the data, and wrote the manuscript; X.L., K.L., L.H., W.P., W.W., S.Z., and H.Z. bred the mice and performed experiments; K.O.L. edited the manuscript and provided intellectual input; B.Z. conceived, supervised, and organized the study.

## Data availability

All data included in this study are included in the manuscript. The raw data are available upon request.

