## Supplementary Figures 1 to 9 for "The robust, high-throughput, and temporally regulated roxCre and loxCre reporting systems for genetic modifications *in vivo*"

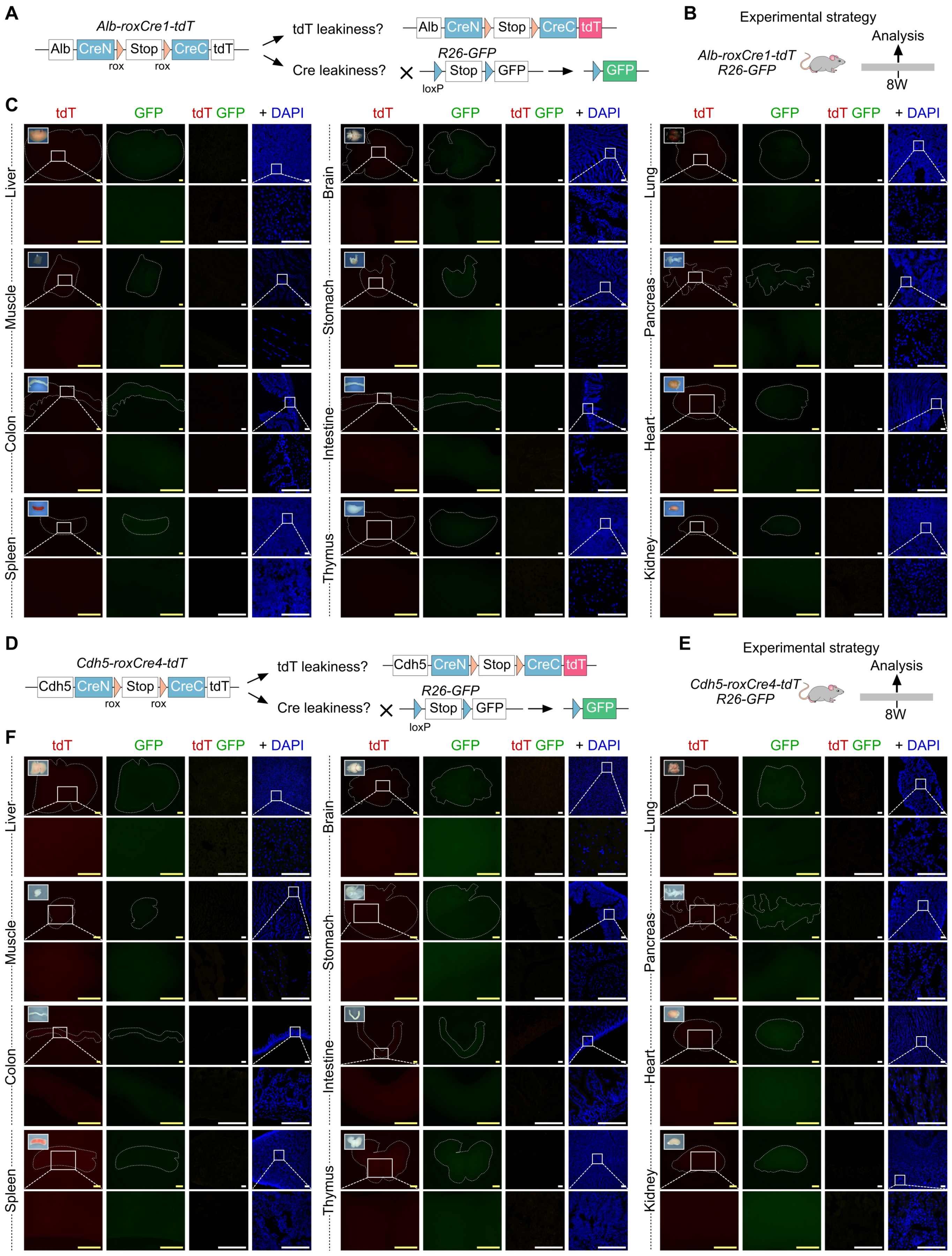


**Supplementary Figure 1. No leakiness in *Alb-roxCre1-tdT;R26-GFP* and *Cdh5-roxCre4-tdT;R26-GFP* mice.**
**A**, **B**, **D**, and **E.** Schematics showing the experimental strategies for examining leakiness of Cre and fluorescence reporters. **C** and **F.** Whole-mount fluorescence and immunostaining sections of *Alb-rox-Cre1-tdT;R26-GFP* and *Cdh5-roxCre4-tdT;R26-GFP* mice. Inserts are bright-field images. Scale bars, yellow, 1 mm; white, 100 μm. Each image is representative of 5 individual biological samples.

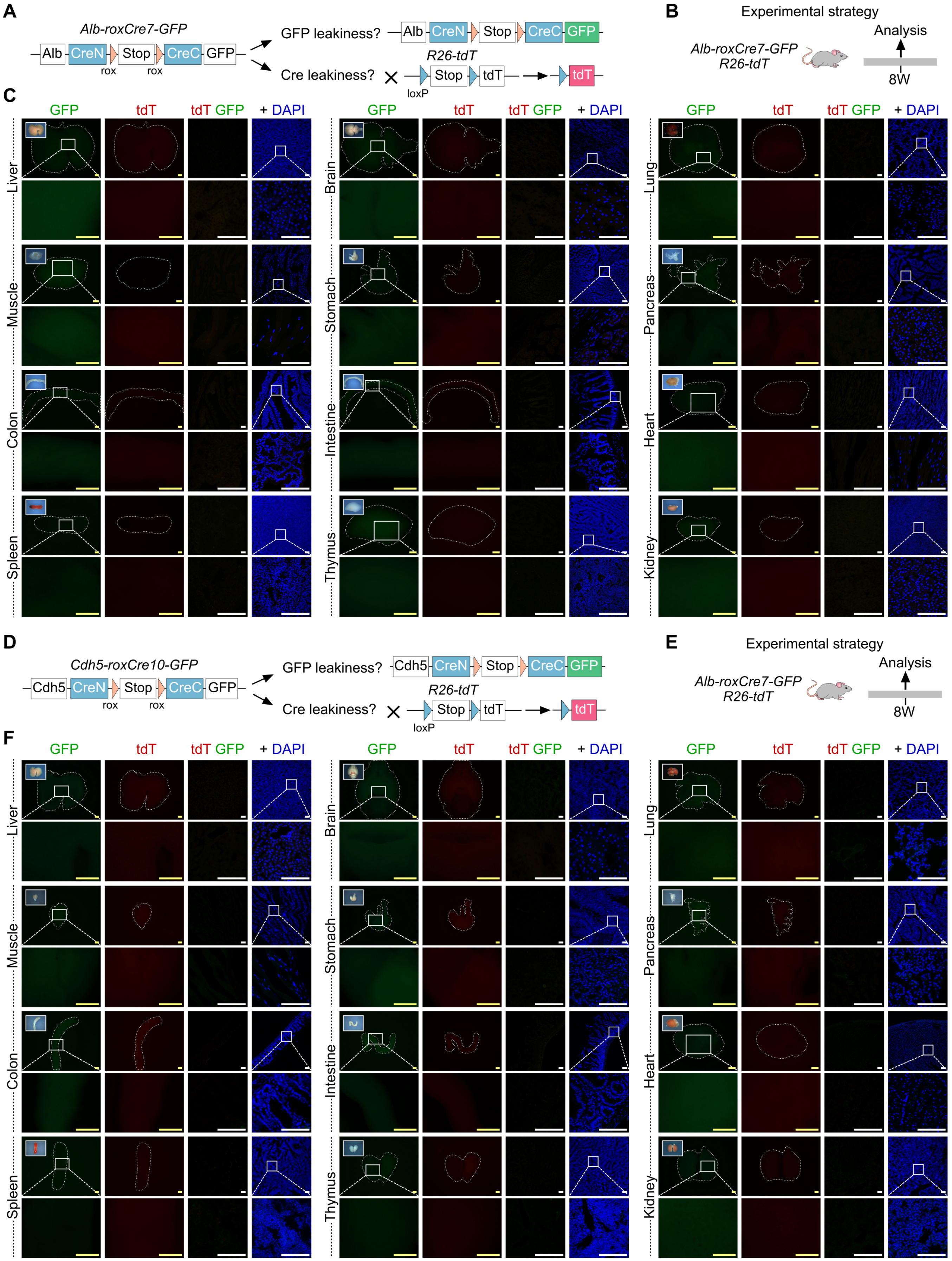


**Supplementary Figure 2. No leakiness in *Alb-roxCre7-GFP;R26-tdT* and *Cdh5-roxCre10-GFP;R26-tdT* mice.**
**A**, **B**, **D**, and **E.** Schematics showing the experimental strategies for examining leakiness of Cre and fluorescence reporters. **C** and **F**. Whole-mount fluorescence and immunostaining sections of *Alb-rox-Cre7-GFP;R26-tdT* and *Cdh5-roxCre10-GFP;R26-tdT* mice. Inserts are bright-field images. Scale bars, yellow, 1 mm; white, 100 μm. Each image is representative of 5 individual biological samples.

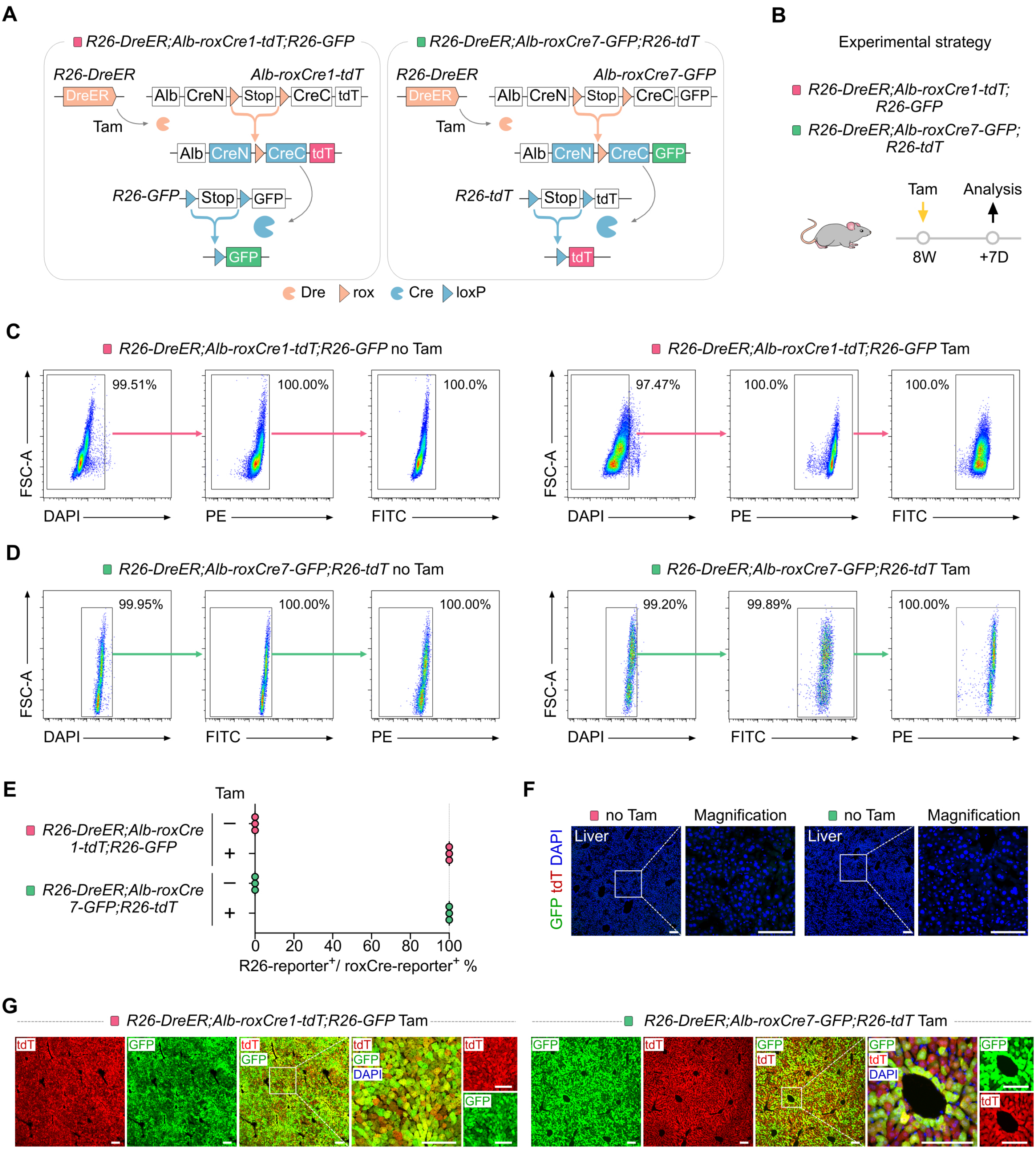


**Supplementary Figure 3. The mCre-loxP recombination efficiency can't be characterized by easily-recombinased reporter.**
**A.** The design of mice mating. **B.** The experimental strategy. **C** and **D.**Gating strategy for analyzing the ratio of R26-reporter$^+$ heps. **E.** The dot plots shows the ratio of R26-reporter$^+$ heps in mCre1-tdT$^+$ heps is equivalent to mCre7-GFP$^+$ heps. **F.** Immunostaining for GFP and tdT on *R26-DreER;Alb-roxCre1-tdT;R26-GFP* and *R26-DreER;Alb-roxCre7-GFP;R26-tdT* no Tam injection liver sections. **G.** Immunostaining for GFP and tdT on Tam-induced *R26-DreER;Alb-roxCre1-tdT;R26-GFP* and *R26-DreER;Alb-roxCre7-GFP;R26-tdT* liver sections. Scale bars, white, 100 µm. Each image is representative of 5 individual biological samples.

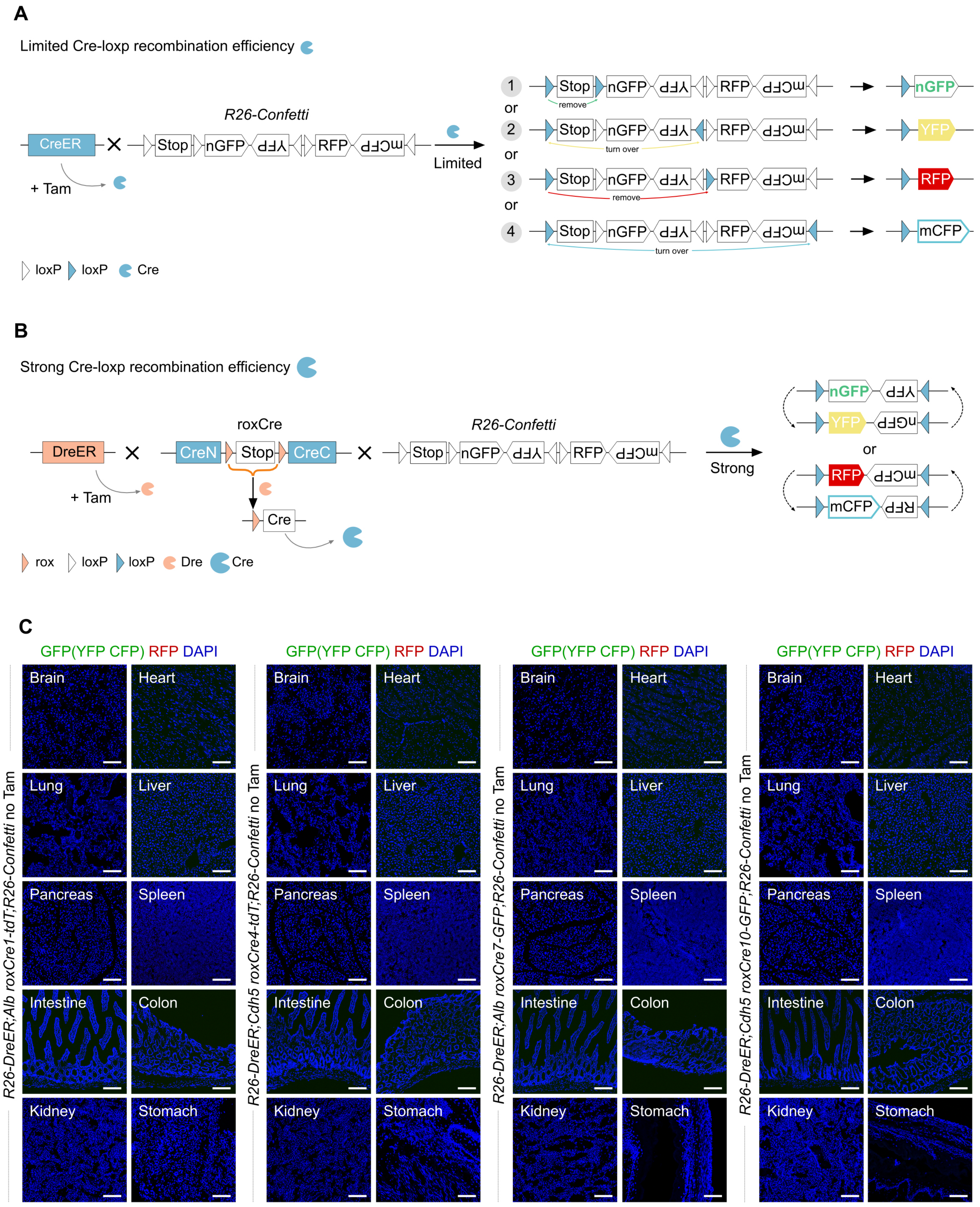


**Supplementary Figure 4. Cre-loxP recombination among different mice.**
**A.** CreER mouse lines which own limited recombination efficiency would recombine *R26-Confetti* into YFP, or nGFP(nuclear GFP), or mCFP(membrane CFP), or RFP. **B.** Tam-induced Dre-rox recombination yielded *mCre* mice can show stronger recombination efficiency and continuously turn over the sequence in the middle of two loxp sites that have the opposite transcription direction. **C.** Immunostaining for GFP(the GFP antibody can also recognize YFP and CFP protein), RFP, and DAPI on *R26-DreER;Alb-roxCre1-tdT;R26-Confetti*, *R26-DreER;Cdh5-roxCre4-tdT;R26-Confetti*, *R26-DreER;Alb-roxCre7-GFP;R26-Confetti*, and *R26-DreER;Cdh5-roxCre10-GFP;R26-Confetti* no Tam injection organs' sections. Scale bars, white, 100 μm. Each image is representative of 5 individual biological samples.

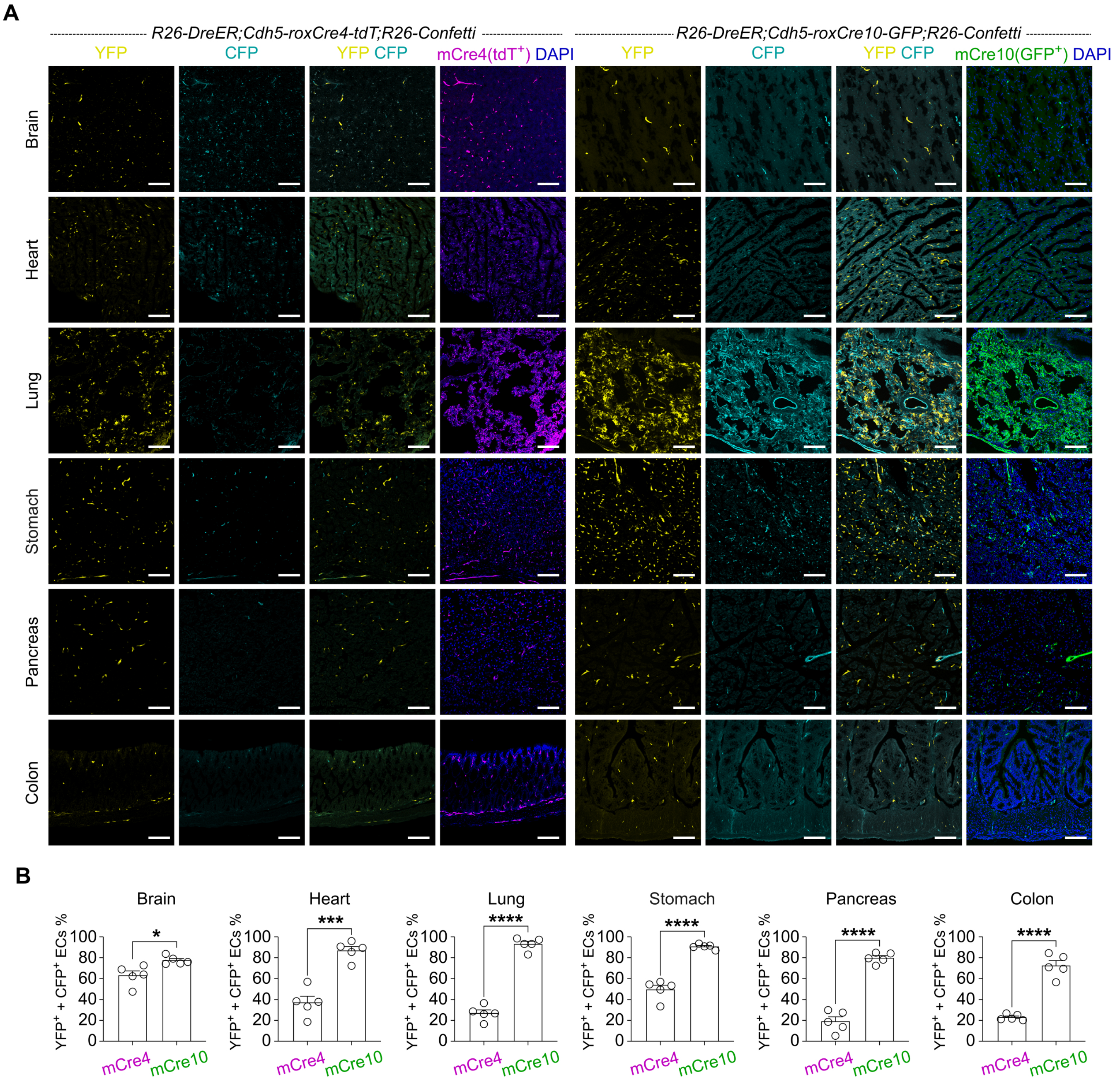


**Supplementary Figure 5. Comparison of mCre recombination among mCre4 and mCre10.**
**A.** Immunostaining for YFP and CFP on tissue sections of *R26-DreER;Cdh5-roxCre4-tdT;R26-Confetti* and *R26-DreER;Cdh5-roxCre10-GFP;R26-Confetti* mice. **B.** Quantification of the percentage of ECs expressing YFP and/or CFP. Data are the means ± SEM; n = 5. *$P$ < 0.05, ***$P$ < 0.001, ****$P$ < 0.0001 by student's $t$ test. Scale bars, white, 100 μm. Each image is representative of 5 individual biological samples.

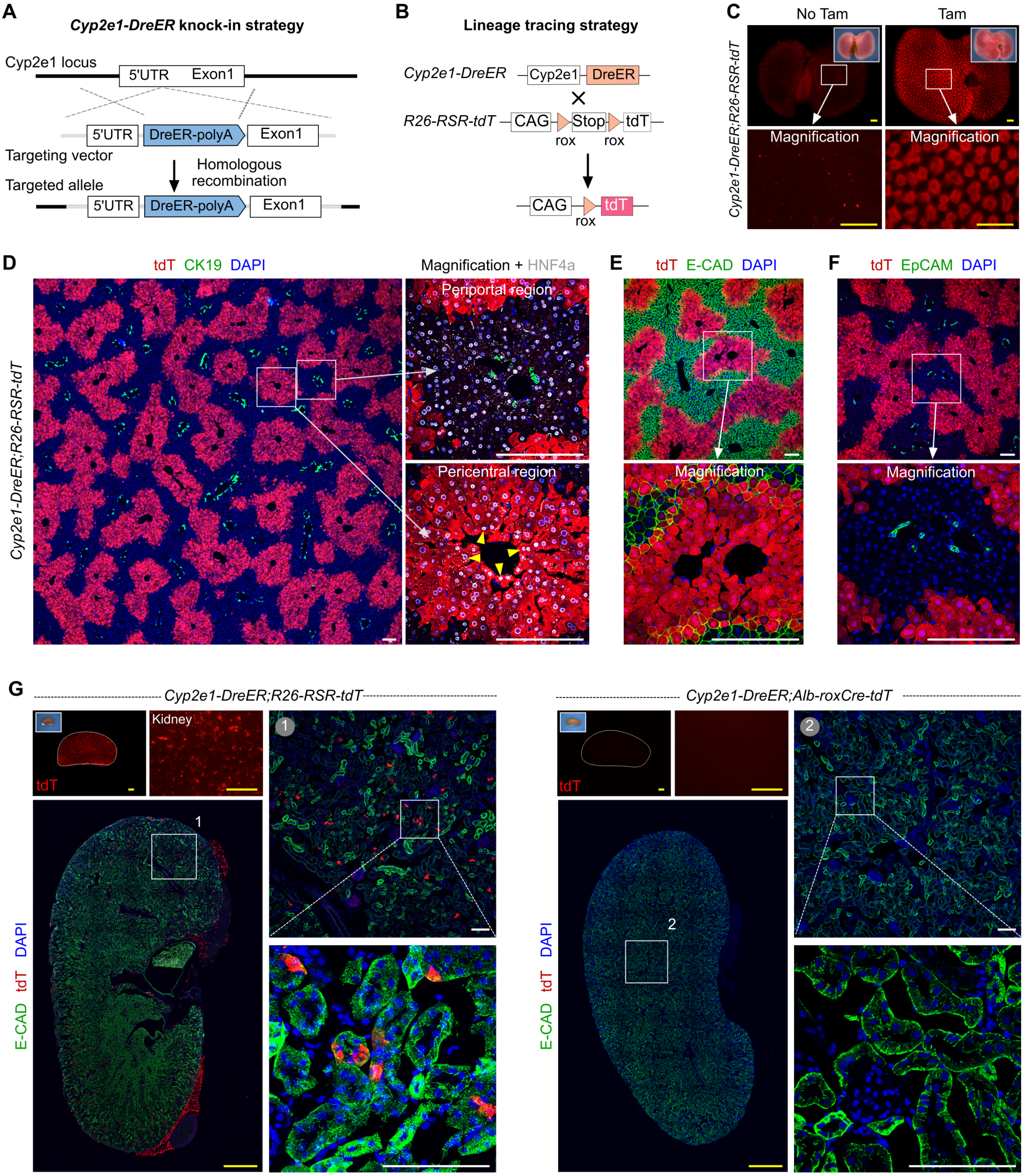


**Supplementary Figure 6. Generation and characterization of *Cyp2e1-DreER* mouse.**
**A.** A schematic showing the knock-in strategy for generation of *Cyp2e1-DreER* allele using CRISPR/Cas9. **B.** A schematic showing the genetic lineage tracing strategy. **C.** Whole-mount fluorescence images of livers from *Cyp2e1-DreER;R26-RSR-tdT* mice treated with or without Tam. Inserts are bright-field images. **D.** Immunostaining for tdT, CK19, and HNF4a on liver sections. Yellow arrowheads, CYP2E1$^+$HNF4a$^+$ hepatocytes. **E.** Immunostaining for tdT and the periportal hepatocyte marker E-CAD shows most CYP2E1$^+$ hepatocytes are located in the peri-central region but not the peri-portal region. **F.** Immunostaining for tdT and the epithelial cells markers EpCAM shows CYP2E1$^+$ cells are not biliary epithelial cells. **G.** Whole-mount fluorescence and immunostained sections of kidneys from *Cyp2e1-DreER;R26-RSR-tdT* (left panel) and *Cyp2e1-DreER;Alb-roxCre-tdT* (right panel) mice. Scale bars, yellow 1 mm; white 100 μm. Each image is representative of 5 individual biological samples.

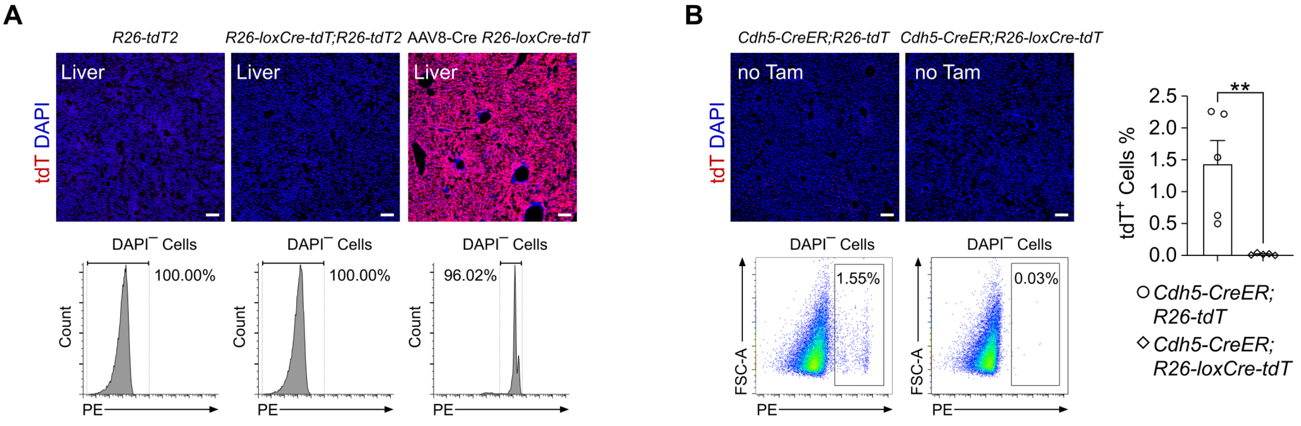


**Supplementary Figure 7. The characterization of *R26-loxCre-tdT* mouse line.**
**A.** *R26-loxCre-tdT* is not leaky when crossed with *R26-tdT2*. **B.** As for some leaky *CreER* mouse lines, such as *Cdh5-CreER*, *R26-loxCre-tdT* can limit CreER's leakiness. Data are the means ± SEM; n = 5 mice. $^{**}P < 0.01$ by student's *t* test. Scale bars, white 100 μm. Each image is representative of 5 individual biological samples.

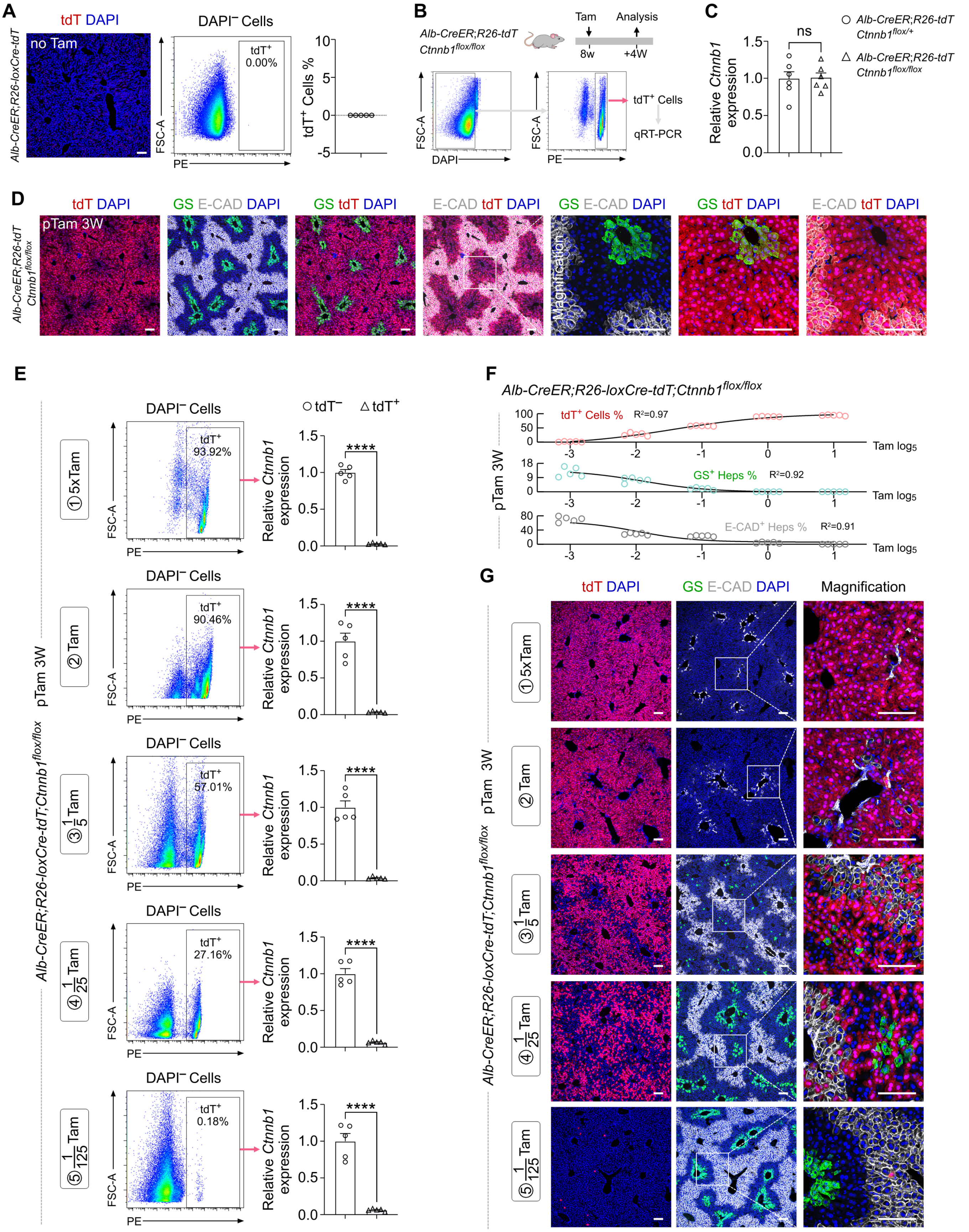

**Supplementary Figure 8. *Ctnnb1* is knocked-out in tdT$^+$ Heps of *Alb-CreER;R26-loxCre-tdT* mouse line.**

**A.** *Alb-CreER;R26-loxCre-tdT* is no leaky. **B.** Sorting out tdT$^+$ Heps of *Alb-CreER;R26-tdT;Ctnnb1$^{flox/flox}$* on the 7th day post one dose of tam injection. **C.** The relative *Ctnnb1* expression by qRT-PCR. **D** and **G.** Immunostaining for tdT, GS, E-CAD, and DAPI on liver sections. **E.** Sorting out tdT$^+$ Heps and tdT$^-$ Cells of *Alb-CreER;R26-tdT;Ctnnb1$^{flox/flox}$* after 3 weeks(W) post concentration-gradient tam injection for the relative *Ctnnb1* expression qRT-PCR. Data are the means ± SEM; n = 5. ****$P$ < 0.0001 by student's $t$ test. **F.** The dose-response curve of the tdT$^+$ hepatocytes ratio by FASC (top), the GS$^+$ hepatocytes ratio by staining(middle), and the E-CAD$^+$ hepatocytes ratio by staining(bottom) with $\log_5$ of the Tam concentration. Scale bars, white 100 μm. Each image is representative of 5 individual biological samples.

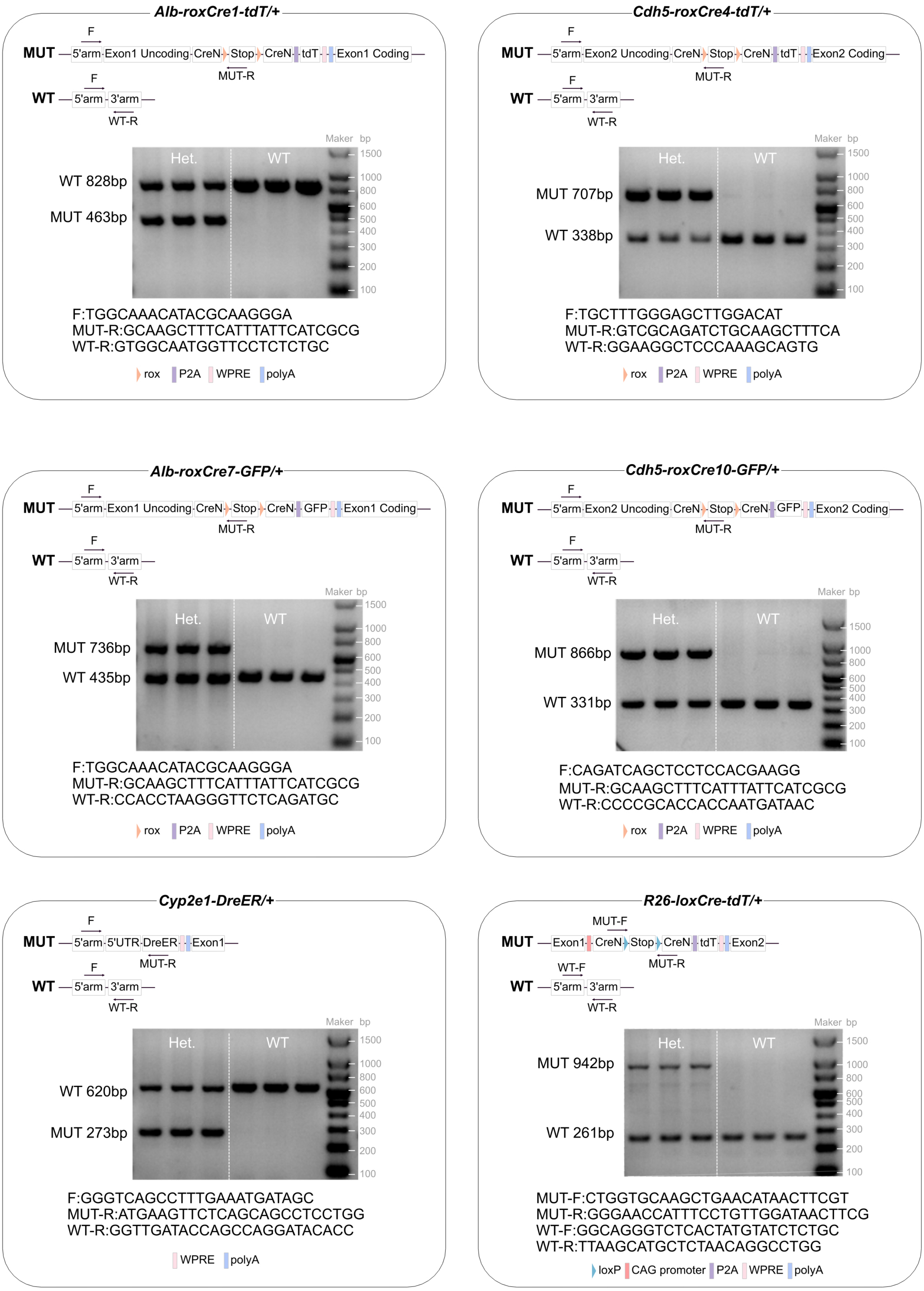


**Supplementary Figure 9. Genotyping information for new mouse lines in this study.**
The mouse line design, primer design, representative results of PCR genotyping, and primer sequences for six mouse lines in this study.
